# Local PI(4,5)P_2_ pool dynamics detected by the coincidence biosensor tubbyCT maintain the integrity of ER-PM junctions during PLC signaling

**DOI:** 10.1101/2020.09.25.313403

**Authors:** Veronika Thallmair, Lea Schultz, Saskia Evers, Christian Goecke, Sebastian Thallmair, Michael G. Leitner, Dominik Oliver

## Abstract

Phosphoinositides (PIs) are important signaling molecules and determinants of membrane identity in the eukaryotic plasma membrane, where they multi-task in divergent signaling pathways. Signaling pleiotropy likely depends on distinct PI pools in the same membrane, although the physical definition of such pools has remained ambiguous. PI(4,5)P_2_, specifically, is also the precursor for the second messengers in the Gq/PLC pathway, IP_3_ and DAG, and is broken down by PLCβ during signaling. Endoplasmic reticulum-plasma membrane contact sites (ER-PM junctions) have emerged as central hubs for lipid transport between both membranes, and specifically for PI homeostasis by supplying the PM with phosphatidylinositol.

Here we show that the tubby protein, by virtue of its C-terminal tubby-domain, preferentially localizes to ER-PM junctions by binding to both PI(4,5)P_2_ and the ER-PM tether E-Syt3. Under conditions of vigorous PI(4,5)P_2_ consumption by PLCβ, additional recruitment of tubby revealed an increase of a local PI(4,5)P_2_ pool fed by local synthesis through PI kinases. Inhibition of this pool-filling process led to the release of the ER-PM tethers, E-Syts, from the membrane and hence to loss of integrity of the ER-PM contact sites.

We conclude that spatiotemporal metabolic channeling of PI synthesis initiated by non-vesicular transport in the ER-PM junctions specifies a local pool of PI(4,5)P_2_ that is pivotal for the maintenance of homeostatic functions during global depletion of PI(4,5)P_2_. The findings further suggest that the tubby-like proteins (TULPs), so far known to impact on energy homeostasis and obesity through primary cilia signaling, have an additional function at ER-PM junctions.

**HIGHLIGHTS:**

- The tubby domain preferentially assembles into ER-PM junctions due to coincidence detection of PI(4,5)P_2_ and E-Syt3
- Tubby recruitment reveals an increase of a local pool of PI(4,5)P_2_ in ER-PM junctions during PLCβ signaling
- Junctional PI(4,5)P_2_ dynamics require local synthesis of PI(4,5)P_2_
- Local PI(4,5)P_2_ supply is required for integrity of ER-PM junctions during PLCβ activity.

## INTRODUCTION

Phosphoinositides (PIs) are minor components of all eukaryotic membranes that nevertheless are pivotal in orchestrating many cell functions. PIs are derived by differential phosphorylation of phosphatidylinositol (PtdIns) through complex metabolic pathways involving a multitude of PI kinases and phosphatases (Balla, 2013).

PIs can act as instructive signals (i.e., classical messengers), that through concentration dynamics can switch on or off distinct signaling pathways (Balla, 2013). Moreover, PIs provide a pivotal membrane identity code, in that different membrane compartments are characterized by specific patterns of PI abundance, which in turn are read by highly specific PI-binding protein modules (Balla, 2013; Di Paolo and De Camilli, 2006; Hammond and Balla, 2015) and less specific electrostatic interactions mediated by polybasic motifs present in many membrane-interacting proteins (Hammond et al., 2012; Heo et al., 2006). The eukaryotic plasma membrane (PM) is characterized by the co-abundance of two PIs, phosphatidylinositol-4-phosphate (PI(4)P) and phosphatidylinositol-4,5-bisphosphate (PI(4,5)P_2_) (Balla, 2013). Importantly, PI(4,5)P_2_ also serves an additional role in signaling by providing the substrate for the hydrolytic generation of the second messengers IP_3_ and diacylglycerol (DAG) by phospholipase C (PLC). Particularly, G-protein signaling through numerous Gq/11-coupled receptors engages PLCβ isoforms. Under such conditions, the PM pool of PI(4,5)P_2_ can be turned over rapidly, necessitating resynthesis from its precursor, phosphatidylinositol (PtdIns). However, PtdIns is generated in the endoplasmic reticulum (ER) and thus needs to be transported to the PM for further phosphorylation by PI kinases.

ER-PM junctions are sites of close apposition of the ER with the PM, mediated by various types of tether proteins including the extended synaptotagmins (E-Syt1-3; (Saheki and De Camilli, 2017a). These membrane contact sites recently emerged as a central hub for membrane lipid homeostasis by mediating transport of phospholipids and sterols between the ER and the PM (Pemberton et al., 2019; Saheki and De Camilli, 2017a). In terms of phosphoinositide lipids, their precursor, PtdIns, is delivered from the ER to the PM by non-vesicular transport mechanisms at ER-PM junctions. Specifically, the Nir2/3 proteins mediate counter-transport of PtdIns against phosphatidic acid that accumulates in the PM during PLC signaling (Chang and Liou, 2015; Kim et al., 2015). Additionally, TMEM24 has been identified as a PtdIns transfer protein at these contact structures (Lees et al., 2017a). Vice versa, E-Syt1 mediates DAG clearance towards the ER (Saheki et al., 2016). During Gq/PLC signaling, recruitment and enhanced PM binding of tether proteins results in tightening of the inter-membrane cleft and expansion of the contact area (Chang et al., 2013; Fernández-Busnadiego et al., 2015; Giordano et al., 2013), and the delivery of PtdIns to the PM is upregulated by recruitment of the PtdIns transport proteins, Nir2 and TMEM24 into the junctions (Chang et al., 2013; Kim et al., 2013; Lees et al., 2017a).

Paradoxically, this transport machinery refilling the PI(4,5)P_2_ pool of the PM critically depends on PI(4,5)P_2_ to work. Most of the tethering proteins that establish the connection between both membranes are ER-resident membrane proteins that anchor to the PM by binding to PI(4,5)P_2_ and sometimes other anionic lipids (Besprozvannaya et al., 2018; Giordano et al., 2013; Sohn et al., 2018). Indeed, experimental depletion of PI(4,5)P_2_ disrupted E-Syt-mediated ER-PM junctions (Giordano et al., 2013). So how can lipid transfer in the ER-PM junctions be kept up when it is needed most - during strong depletion of PI(4,5)P_2_ by PLC signaling?

One possibility for enabling continued functionality of ER-PM junctions during PLC signaling is that the pool of PI(4,5)P_2_ required for tether binding to the PM at ER-PM junctions is distinct from the PI(4,5)P_2_ accessible to PLC-mediated hydrolysis. Thus, the existence of PI(4,5)P_2_-enriched microdomains at ER-PM junctions was proposed (Maléth et al., 2014). However, direct experimental evidence for such spatial enrichment is still lacking (reviewed e.g., by (Chang and Liou, 2016).

More generally, the highly diverse yet specific tasks of PI(4,5)P_2_ (and PI(4)P) in the PM may be supported by the existence of various distinct pools of these lipids. Such pools may be organized spatially, forming PI(4,5)P_2_ (micro)domains generated either by locally confined synthesis as it has already been shown in specific processes such as phagocytosis (Levin et al., 2015) and endocytosis (He et al., 2017), or by electrostatic clustering induced by membrane-associated proteins (McLaughlin and Murray, 2005). On the other hand, there is evidence for distinct functional pools of PI(4,5)P_2_ and PI(4)P in the PM without an obvious spatial organization of these populations (Hammond et al., 2012; Hammond et al., 2009).

An additional layer of complexity in determining the specificity of phosphoinositide signaling is suggested by the concept of instructive regulation of phosphoinositide kinases by context-specific delivery of the substrate, PtdIns, through phosphatidylinositol transfer proteins (PITPs) (Grabon et al., 2015). Thus, in yeast, engagement of the various Sec14-homologous PITPs specifies the biological activity of PI(4)P synthesized by the same PI4K enzyme (Grabon et al., 2015). It is intriguing to speculate that in animal cells such metabolic channeling may occur at ER-PM junctions to generate functionally and even spatially distinct populations of PI lipids. However, it has so far remained unknown if and how the transport of PtdIns by the lipid transfer proteins Nir2/3 or TMEM24 at ER-PM junctions is coordinated with the enzymatic phosphorylation steps at the PM that generate the major PIs, i.e. PI(4)P and PI(4,5)P_2_ (Pemberton et al., 2019).

Here we identify the C-terminal domain of the tubby protein (tubbyCT) as a PI(4,5)P_2_ sensor selectively enriched at ER-PM junctions by interaction with E-Syt3 and PI(4,5)P_2_. This co-incidence detection property allowed for monitoring of local pools of PI(4,5)P_2_ in these domains. Unexpectedly, Gq-coupled receptor activation of PLC signaling induced an increase of the local PI(4,5)P_2_ concentration that was detected as recruitment of tubbyCT, while concomitantly PI(4,5)P_2_ was depleted from the overall membrane. This inverse kinetic behavior identifies a local pool of PI(4,5)P_2_ that escaped detection by conventional PI(4,5)P_2_ biosensors. Blocking the buildup of this local PI(4,5)P_2_ transient disrupted tethering of ER-PM junctions. Thus the local PI(4,5)P_2_ pool serves to maintain homoeostatic function of the contact sites by securing tethering of both membranes, while allowing the discharge of distinct PI(4,5)P_2_ dynamics in the bulk membrane. The local PI(4,5)P_2_ source may result from the spatially focused delivery of PtdIns to the junctional PM, further suggesting spatial metabolic channeling of PtdIns to a PI(4,5)P_2_-synthetic machinery localized to ER-PM junctions.

## RESULTS

### TubbyCT associates with the PM during PLC-mediated consumption of PI(4,5)P_2_

When tubbyCT is expressed in various cell types, it associates with the PM as shown by membrane-localized fluorescence of fused GFP (Hackelberg and Oliver, 2018; Halaszovich et al., 2009; Hardie et al., 2015; Nelson et al., 2008; Quinn et al., 2008; Santagata et al., 2001). Membrane association has been demonstrated to be mediated by specific binding to PI(4,5)P_2_ as indicated by a variety of approaches including in-vitro lipid binding assays (Santagata et al., 2001), crystallography (Santagata et al., 2001), molecular dynamics simulations (Thallmair et al., 2020) and manipulation of PI(4,5)P_2_ concentration in living cells (Halaszovich et al., 2009; Mavrantoni et al., 2015; Szentpetery et al., 2009).

However, the lipid binding properties of tubbyCT have remained puzzling. Thus, activity of the Gq/PLCβ pathway, which is generally thought to involve depletion of PI(4,5)P_2_ (at least when strongly activated), readily displaces tubbyCT from the PM, as expected, in some experimental settings (Hackelberg and Oliver, 2018; Santagata et al., 2001). In contrast, other studies show reluctant response to PLCβ activation, particularly when compared to the popular PI(4,5)P_2_ sensor domain PLCΔ1-PH-GFP (Leitner et al., 2019; Quinn et al., 2008; Szentpetery et al., 2009).

Unexpectedly and paradoxically, we observed that activation of Gq/PLCβ by muscarinic M1 receptor (M1R) in CHO cells resulted in recruitment of tubbyCT to the PM as demonstrated by live-cell TIRF microscopy (**Fig. 1A**), which would suggest an increase in PI(4,5)P_2_ concentration rather than the expected decrease. However, the alternative PI(4,5)P_2_ sensors PLCΔ1-PH (Fig. 1A) or ENTH-GFP (Leitner et al., 2019) dissociated from the membrane under identical experimental conditions, attesting to the expected decrease in global PI(4,5)P_2_.

**Figure 1.**
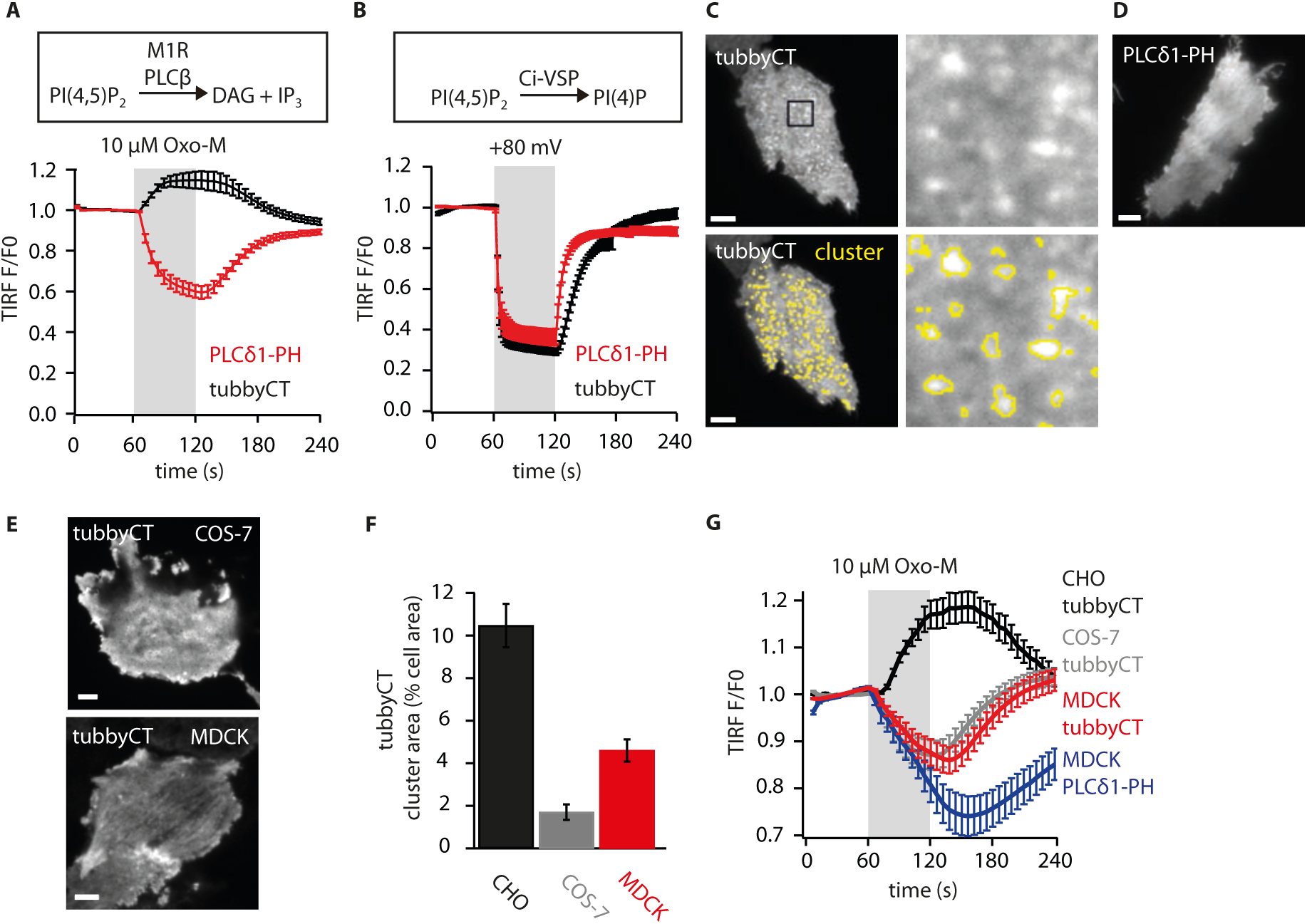
PLC-induced recruitment and microdomain PM organisation of the C-terminal domain of the tubby protein. **(A)** Opposite membrane association dynamics of GFP-tubbyCT and PLCδ1-PH-GFP following activation of PLC-β by stimulation of Gq-coupled receptors. Membrane association of either domain was monitored by TIRF microscopy in CHO cells (n = 32 and 34, respectively) co-transfected with M1 receptors. M1R agonist oxotremorine-M (Oxo-M; 10 µM) was applied as indicated (grey bar). TIRF intensity is plotted normalized to signal before agonist application. **(B)** Dissociation of both domains upon depletion of PI(4,5)P_2_ by the voltage-activated PI(4,5)P_2_ phosphatase Ci-VSP. CHO cells co-expressing either PI(4,5)P_2_-binding domain with Ci-VSP were monitored by TIRF-M as in (A) while membrane potential was switched between -80 mV and +80 mV by whole-cell patch-clamp with timing as indicated by grey bar (tubbyCT, n = 7 cells; PLCδ1-PH, n = 9) **(C)** Representative TIRF image of a CHO shows pronounced spatial clustering of GFP-tubbyCT. Yellow lines in lower panels indicate cluster areas as detected by the sectioning algorithm described in Methods. Right panels shows enlargement of the rectangular region indicated. **(D)** Representative TIRF image of CHO cell expressing PLCδ1-PH-GFP demonstrating lack of clustering. **(E)** Minor clustering of tubbyCT expressed in COS-7 and MDCK cells. TIRF imaging as in (C). **(F)** Cell area occupied by GFP-tubbyCT clusters, quantified from TIRF images by sectioning algorithm as in (C) (CHO, n = 43; COS-7, n = 28; MDCK, n = 31 cells). **(G)** PI(4,5)P_2_ sensor dynamics in different cell types upon activation of PLCβ via co-expressed M1R. CHO, n = 67 cells; COS-7, n = 35; MDCK with tubbyCT, n = 117; MDCK with PLCδ1-PH, n = 26. All data are given as mean ± SEM; Scale bars, 5 µm (C - E).

We therefore first re-evaluated the PI(4,5)P_2_ dependence of membrane association of tubbyCT in CHO cells by depleting PI(4,5)P_2_ with the voltage-activated PI(4,5)P_2_ phosphatase Ci-VSP (Halaszovich et al., 2009; Iwasaki et al., 2008). Activation of co-expressed Ci-VSP by depolarization resulted in rapid and reversible dissociation of tubbyCT from the PM (**Fig. 1B**), confirming PI(4,5)P_2_ as a principal determinant of membrane association. Moreover, PM dissociation was quantitatively similar to that of PLCδ1-PH. Slower re-association during resynthesis of PI(4,5)P_2_ following wash-out of the agonist suggests that its PI(4,5)P_2_ affinity is lower than the affinity of PLCδ1-PH. Similar conclusions were previously drawn from careful titration of PI(4,5)P_2_ (Halaszovich et al., 2009; Leitner et al., 2019), and recent molecular dynamics simulations of PI(4,5)P_2_ binding (Thallmair et al., 2020).

### TubbyCT, but not other PI(4,5)P_2_-binding domains, clusters in PM domains

These paradoxical findings may indicate that tubbyCT associated with a distinct pool of PI(4,5)P_2_ that behaves differently from bulk PI(4,5)P_2_ in the PM in response to PLCβ activity. Such a separate PI(4,5)P_2_ pool may be organized spatially in separate domains. We therefore examined the membrane distribution of tubbyCT by TIRF microscopy. Indeed, GFP-tubbyCT expressed in CHO cells showed a pronounced enrichment in distinct domains, with lower abundance throughout the bulk membrane (**Fig. 1C**). These clusters occupied 10.5 ± 0.9 % of the cell area imaged by TIRF-M with an average density of 0.66 ± 0.03 µm^-2^ (n = 57 cells). Such clustering was not observed with the PI(4,5)P_2_-binding domains PLCδ1-PH or GFP-ENTH (**Fig. 1D; Suppl. Fig. 1A,B**). We note, though, that a minor enrichment of PLCδ1-PH-GFP was occasionally detectable in domains defined by tubbyCT **(Suppl. Fig. 1A)**. We conclude that tubbyCT selectively bound to a PI(4,5)P_2_ pool not preferentially recognized by PLCδ1-PH or ENTH domains. Also, the general membrane marker lyn11-RFP (Inoue et al., 2005) was distributed homogenously despite pronounced clustering of co-expressed tubbyCT in the same membrane area, demonstrating that local GFP-tubbyCT accumulation is not an imaging artifact resulting from local membrane folding or invagination (**Suppl. Fig.1C**).

Of note, clustering of tubbyCT was much less pronounced in two other cell lines examined (COS-7, MDCK; **Fig. 1E,F**). In these cells, activation of the Gq/PLCβ pathway induced dissociation of tubbyCT from the membrane, although to a lower degree than PLCδ1-PH (**Fig. 1G**). Thus, the occurrence of PLCβ-dependent membrane recruitment of tubbyCT correlated with the clustering into spatial domains.

### Dynamic recruitment of tubbyCT into local domains

We next explored the spatiotemporal properties of tubbyCT dynamics by TIRF imaging. Pixel-wise analysis of fluorescence intensity changes following receptor stimulation revealed a spatially highly inhomogeneous distribution of tubbyCT binding and unbinding, with discrete punctate foci of strong membrane recruitment, but dissociation of tubbyCT from the membrane throughout larger more contiguous areas (**Fig. 2A**). Typically, foci of tubbyCT increase were found preferentially in the center of the cells, and loss of membrane association occurred more in peripheral regions of the cell.

**Figure 2.**
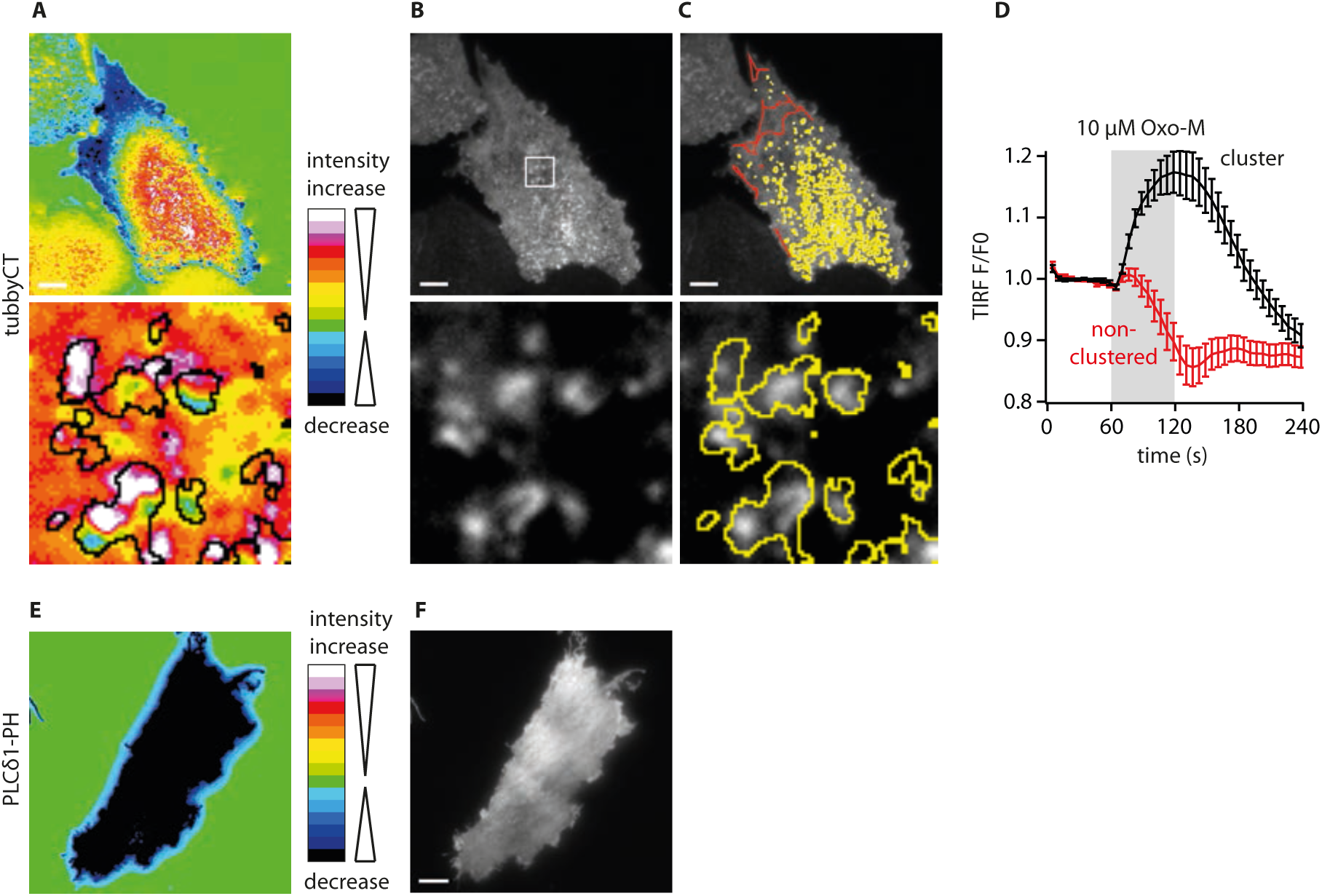
PLC-induced recruitment occurs at tubbyCT-enriched domains. **(A)** Spatial distribution of PLC-induced GFP-tubbyCT membrane dynamics measured by TIRF-M. Shown are color-coded absolute intensity changes induced by 1 min application of Oxo-M (10 µM; co-expression of M1R), calculated pixel-wise. Lower panel, enlarged region (5 µm x 5 µm) as indicated in (B). Black lines indicate calculated borders of the tubby-CT clusters prior to activation of M1R/PLC as shown in (C). **(B)** TIRF image showing the resting GFP-tubbyCT distribution in the same CHO cell. **(C)** Masks delineating regions of enriched (clustered, yellow) and uniform (non-clustered, red) regions as derived with sectioning algorithm (see Methods). **(D)** Time course of GFP-tubbyCT membrane association upon PLCβ activation by stimulation of co-expressed M1R in the clustered (black) and non-clustered (red) regions as defined by the sectioning algorithm (n = 49 cells). **(E,F)** Representative recordings of spatial dynamics following stimulation of M1R (E) and resting distribution of PLCδ1-PH-GFP (F) demonstrate homogenous dissociation from the membrane during PLC signalling. Scale bars, 5 µm.

Since this pattern of tubbyCT dynamics conspicuously resembled the resting distribution of tubbyCT (**Fig. 2B**), we analyzed the spatial relation of tubbyCT dynamics to the clustering pattern. To this end, we devised a segmentation algorithm to define the domains enriched in tubbyCT (**Fig. 2C**; for details, see Methods). As shown in the lower panel of **Fig. 2A**, regions of high tubbyCT recruitment were largely coincident with the clusters. Consequently, when averaging the dynamics from tubbyCT clusters, strong recruitment in response to receptor stimulation was observed (**Fig. 2D**). In contrast, averaging fluorescence changes of non-cluster areas revealed dissociation of tubbyCT from the membrane, which is similar to the general behavior of other PI(4,5)P_2_ sensors but is apparently masked when averaging over the entire membrane area as done in **Fig. 1A**.

In strong contrast to compartmentalized tubbyCT dynamics, spatial analysis of PLCδ1-PH dynamics shows homogenous dissociation over the entire membrane area (**Fig 2E,F**).

Provided that tubbyCT membrane association results from binding to PI(4,5)P_2_, the tubbyCT clusters therefore demarcate areas of increasing PI(4,5)P_2_ abundance rather than PI(4,5)P_2_ consumption in response to PLCβ activation.

### TubbyCT associates to ER-PM junctions in an E-Syt3-dependent manner

ER-PM junctions play a central role in PI(4,5)P_2_ replenishment following PLCβ activation. Non-vesicular transport of the precursor of PIs, PtdIns, from the ER to the PM is mediated by these structures, and this transport is stimulated by PLC signaling (Chang et al., 2013; Chang and Liou, 2015; Kim et al., 2015).

We thus reasoned that tubbyCT clusters may be related to these membrane contact sites. Typical components of ER-PM junctions are the extended synaptotagmins (E-Syt1-3). E-Syts are molecular tethers connecting the ER with the PM, being anchored in the ER membrane with an N-terminal hairpin loop and binding the PM with C-terminal C2 domains dependent on PI(4,5)P_2_ and Ca^2+^ (Giordano et al., 2013; Saheki and De Camilli, 2017b).

Upon co-expression of tubbyCT with E-Syt1 or E-Syt2, both the fluorescently tagged E-Syt and tubbyCT formed similar patterns of membrane-localized domains, which however showed no substantial spatial correlation (**Figs. 3A,B**). However, tubbyCT co-clustered with E-Syt3-enriched domains, as shown in **Fig. 3C**. Co-localization was also quantitatively confirmed by calculating Pearson’s coefficient for pixel-wise correlation of fluorescence intensities (**Fig. 3F**). Thus, the coefficient was close to zero for correlation of tubbyCT localization with E-Syt1 and E-Syt2, but about 0.3 for E-Syt3. To validate this quantification, we attempted to calibrate the Pearsons’s coefficient by examining experimental conditions where maximal co-localization is expected. Co-expression of GFP-tubbyCT with RFP-tubbyCT or of two E-Syt isoforms known to form heteromers (Giordano et al., 2013), yielded even higher correlation coefficients of around 0.8 (**Fig. 3F, Suppl. Fig. S2**). A more moderate correlation of tubbyCT with E-Syt was expected since visual inspection already reveals a substantial fraction of membrane-localized tubbyCT outside of the tubbyCT clusters that co-localize with E-Syt3. In contrast to tubbyCT, PLCδ1-PH did not substantially co-localize with any of the three E-Syt isoforms (**Suppl. Fig. S3**), although the correlation coefficient was slightly but consistently above zero, possibly refelecting the homogenous distribution spanning the contact sites or resulting from a faint co-enrichment also observed in co-expression with tubbyCT (**Fig. S1A**).

**Figure 3.**
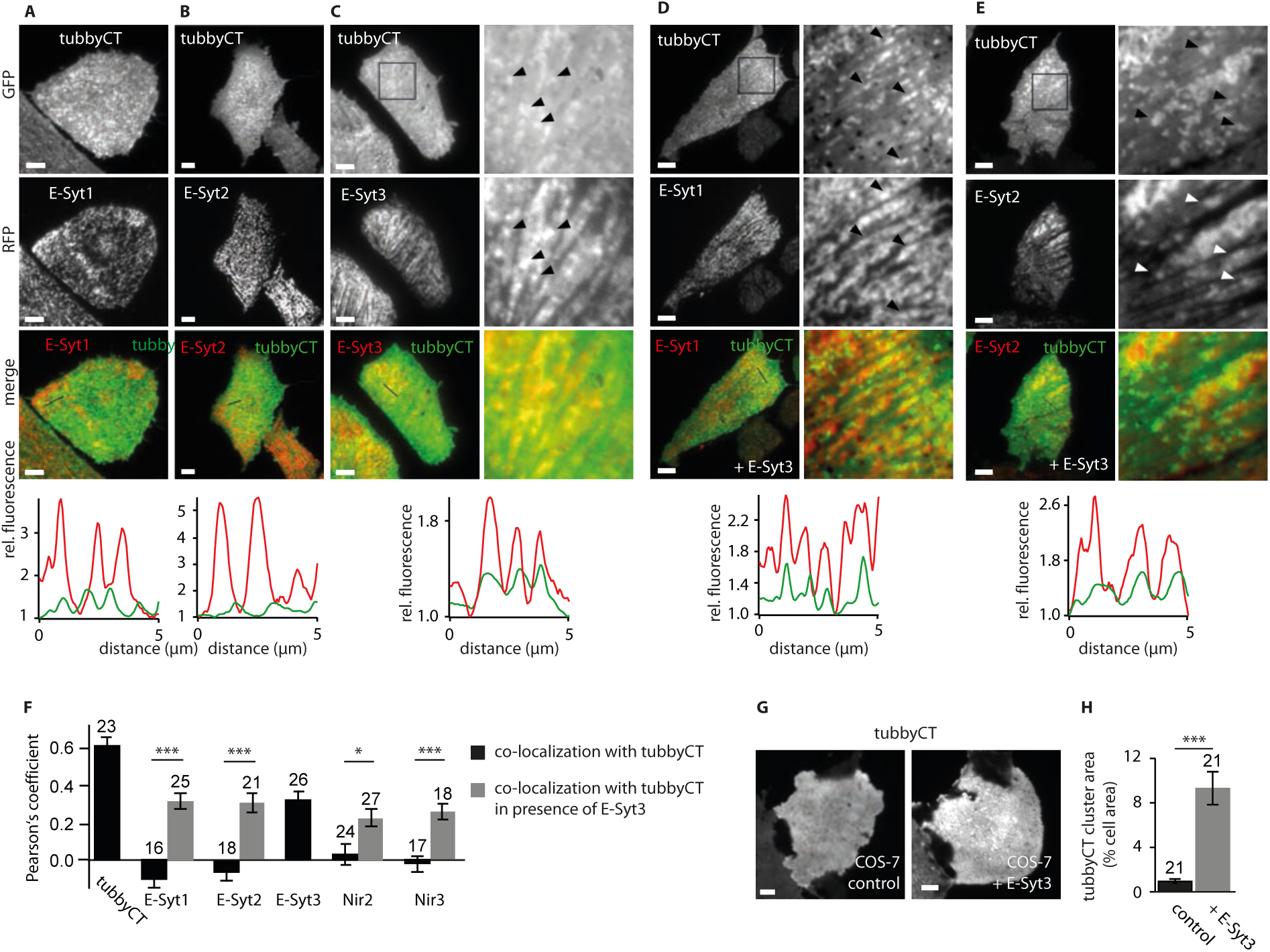
TubbyCT localizes to ER-PM junctions. **(A-C)** Co-localization analysis of GFP-tubbyCT with RFP-E-Syt1, RFP-E-Syt2, and RFP-E-Syt3, respectively. Transiently co-transfected CHO cells were imaged using TIRF microscopy. Lower panels, fluorescence intensity line profiles for representative regions indicated in merged fluorescence images above. **(D, E)** GFP-tubbyCT co-localized with RFP-E-Syt1 (D) and RFP-E-Syt2 (E), when additionally CFP-E-Syt3 was co-overexpressed. **(F)** Pearson coefficients r (mean ± SEM) from cells as in (A-E) and from co-localization analysis of GFP-tubbyCT with RFP-tubbyCT, Nir2-mcherry and Nir3-mcherry, respectively. Pearson’s coefficients obtained from cells expressing GFP-tubbyCT with RFP-tagged proteins only are shown in black, corresponding values from cells additionally expressing CFP-E-Syt3 are shown in grey. Mean ± SEM, number of cells analyzed as indicated. **(G)** In COS-7 cells E-Syt3 over-expression evoked tubbyCT cluster formation (right TIRF image; GFP-tubbyCT + RFP-E-Syt3) not observed when tubbyCT was expressed together with free RFP (control; left). **(H)** Cell area occupied by tubbyCT clusters analyzed from images as in (G); mean ± SEM, numbers of cells indicated; Student’s t test: p = 0.000012). Scale bars, 5 µm.

Given the colocalization with the ER-PM junction protein E-Syt3, we asked whether tubbyCT clusters in fact correspond to ER-PM junctions. Although lacking co-localization with co-expressed ER-PM tethers E-Syt1 and E-Syt2, upon additional over-expression of E-Syt3, tubbyCT also localized to the domains containing either E-Syt1-RFP (**Fig. 3D,F**) or E-Syt2-RFP (**Fig. 3E,F**), i.e. to presumptive ER-PM junctions. We next examined colocalization of tubbyCT with additional junctional proteins. TubbyCT showed no obvious colocalization with the ER-PM components Nir2 and Nir3 (Chang and Liou, 2015; Kim et al., 2013; Kim et al., 2015) when co-expressed with RFP-fusion constructs of these proteins. However, when E-Syt3 was additionally overexpressed, we observed co-localization to domains outlined by either Nir isoform (**Fig. 3F**).

Taken together, we conclude that tubbyCT specifically assembles into ER-PM junctions that contain E-Syt3. This is a unique feature of tubbyCT which was not observed for the other PI(4,5)P_2_-binding domains tested here, namely PLCδ1-PH and ENTH. Accordingly, we assume that the tubbyCT clusters observed in naive CHO cells represent ER-PM junctions containing E-Syt3. Other cells lines, such as COS-7 cells may lack the expression levels of E-Syt3 required to strongly recruit tubbyCT into ER-PM junctions. In fact, when E-Syt3 was over-expressed in COS-7 cells, tubbyCT localization adopted the pronounced clustering characteristic for CHO cells, lending further support to the role of E-Syt3 in localizing tubbyCT into ER-PM junctions (**Fig. 3G,H**).

### Clustering into ER-PM junctions is mediated by coincidence binding to E-Syt3 and PI(4,5)P_2_

Given that E-Syt3 promotes clustering of tubbyCT into ER-PM junctions, we next explored a potential direct interaction between both proteins. Co-IP experiments were performed with CHO cells transfected with myc-tagged tubbyCT and GFP-fused E-Syt constructs. **Fig. 4A** shows that anti-GFP antibodies robustly co-precipitated myc-tubbyCT from cells co-expressing GFP-E-Syt-3, and – to a lesser degree – from cells expressing GFP-E-Syt1 or GFP-E-Syt2. No interaction was observed between E-Syt3 and PLCδ1-PH. These finding indicate complex formation between tubbyCT and E-Syt3, and perhaps a weaker interaction of tubbyCT with E-Syt1 and E-Syt2. However, co-precipitation of tubbyCT with Syt1 and E-Syt2 may as well result from indirect interaction mediated by endogenous E-Syt3, given the known dimerization of E-Syt1/2 with E-Syt3 (Giordano et al., 2013). Thus, E-Syt1/2 likely co-precipitate tubbyCT bound to endogenous E-Syt3. We confirmed co-precipitation of E-Syt3 with E-Syt1 (**Fig. 4A**). Of note, this interpretation is also consistent with the finding that over-expression of E-Syt3 strongly enhances cellular co-localization of tubbyCT with E-Syt1 or E-Syt2 (**Fig.3**).

**Figure 4.**
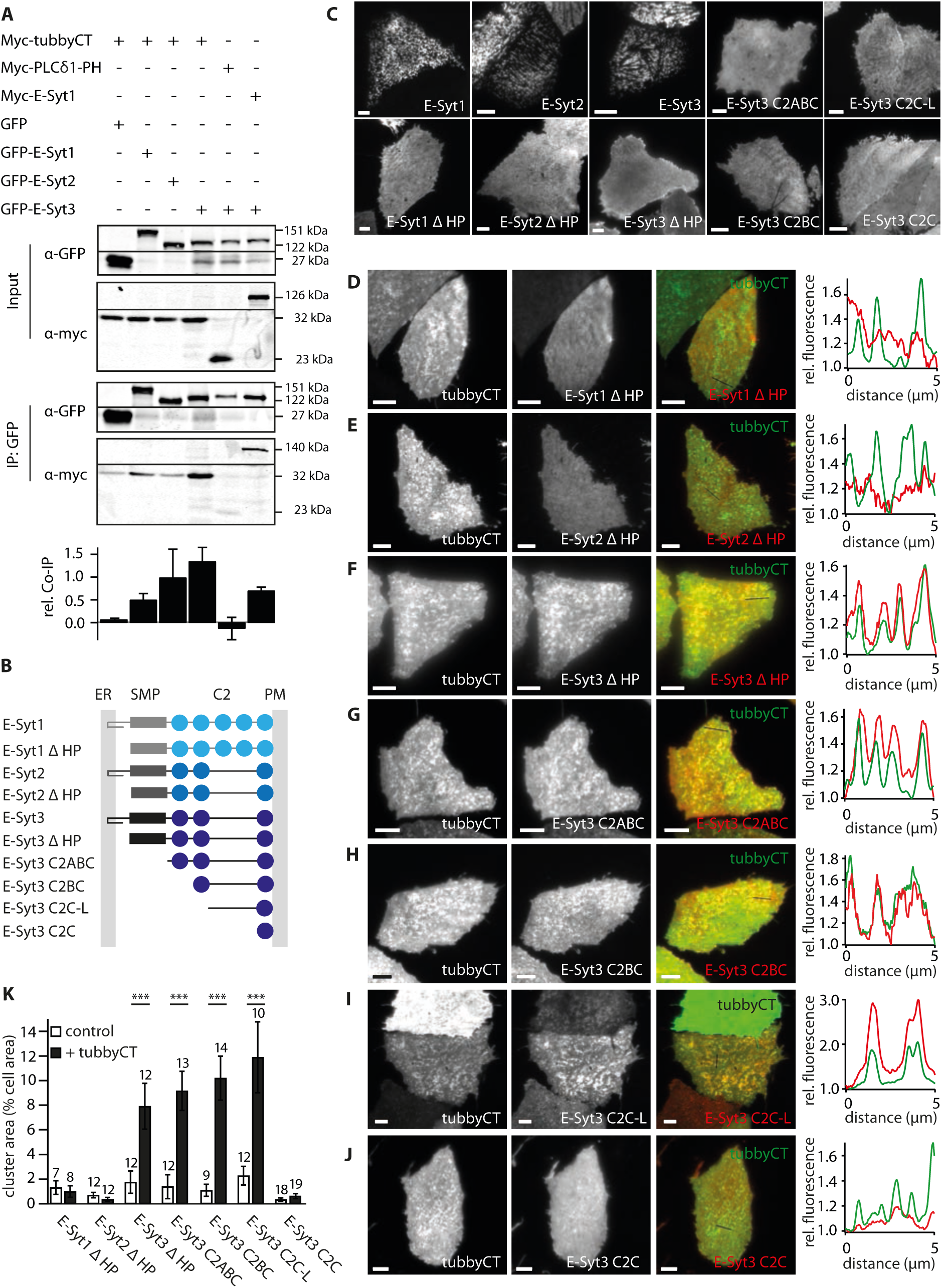
Molecular interaction of tubbyCT with E-Syt3. **(A)**Co-immunoprecipitation of myc-tubbyCT with GFP-tagged E-Syt proteins. CHO cells were transfected with myc- and GFP-tagged constructs as indicated. Protein lysates of cells co-expressing myc-tubbyCT with free GFP and myc-PLCδ1-PH with GFP-E-Syt3 were used as negative controls. Lysates of myc-E-Syt1 and GFP-E-Syt3-expressing cells served as positive control. Representative blots of α-GFP and α-myc stained input and immunoprecipitation are shown. Lower panel shows quantification of four independent experiments (mean ± SEM) (myc-tubbyCT/GFP-E-Syt3 versus myc-tubbyCT/free GFP: p = 0.0250). **(B)**Domain architecture of E-Syt isoforms and E-Syt3 truncation constructs. (SMP domain, grey; C2 domains, blue; HP = ER membrane binding hairpin). **(C)**Representative TIRF images of CHO cells transiently expressing the various RFP-E-Syt constructs shown in (B). All truncation constructs exhibit uniform PM localization. **(D-J)** Representative TIRF images and co-localization analysis in cells co-expressing GFP-tubbyCT with the various E-Syt truncation constructs. Note rescue of clustered distribution and tubbyCT co-localization for E-Syt3 truncation constructs in (F – I). **(K)** Cell area occupied by RFP-E-Syt clusters in presence of free GFP (control) and GFP-tubbyCT analyzed from images as in (D-J). Cluster area was determined by sectioning algorithm described in Methods. (E-Syt3 ΔHP, Student’s t test p = 0.00668; E-Syt3 C2ABC, p = 0.00031; E-Syt3 C2BC, p = 0.00013; E-Syt3 C2C-L, p = 0.00660). Scale bars, 5 µm.

In order to scrutinize such interaction and to localize the site of interaction within the modular domain architecture of E-Syt3, we examined co-localization of tubbyCT with various E-Syt truncation constructs. The E-Syts are anchored in the ER membrane by an N-terminal hairpin structure and bind to the PM with their C-terminal C2 domains (Giordano et al., 2013) as schematically shown in **Fig. 4B**. Accordingly, deletion of the hairpin sequence (E-Syt δHP) in either E-Syt isoform resulted in a homogenous distribution of N-terminally RFP-labeled E-Syt in TIRF imaging (**Fig. 4C**). Thus, association with the ER and consequently with ER-PM junctions was lost (cf. (Idevall-Hagren et al., 2015)), but some association with the PM persisted, likely mediated by the C2 domains.

Co-expression of tubbyCT rescued clustering of E-Syt3-ΔHP, but not E-Syt1-ΔHP or E-Syt2-ΔHP into well-defined domains (**Fig. 4D-F,K**), showing that in a reverse manner, tubbyCT is capable of recruiting E-Syt3 into ER-PM junctions. Also, this finding demonstrates that the tubbyCT/E-Syt3 interaction persists in the absence of the hairpin motif. In a series of further truncations tubbyCT-dependent clustering of truncated E-Syt3 persisted when the SMP domain and additionally the first two (N-terminal) C2 domains (C2A and C2B) were deleted (**Fig.4 G-K**). Thus the minimum construct recruited into clusters by tubbyCT was a C-terminal region comprising the C2B-C2C-linker plus the C2C domain (**Fig.4I**), whereas the isolated C2C domain alone (plus its C-terminal extension) was not sufficient for this interaction with tubbyCT (**Fig.4 J**). Taken together, these experiments localize the interaction site for tubbyCT to the C2B/C2C-linker region of E-Syt3.

While these findings strongly support a direct and isoform-specific interaction of tubbyCT with E-Syt3, the unexpected reciprocal recruitment of E-Syt3-ΔHP by tubbyCT immediately suggested that another important interaction partner additionally accounts for preferential localization of tubbyCT at ER-PM contact sites. Thus we investigated whether increased local PI(4,5)P_2_ concentrations could be involved in tubbyCT clustering. Acute depletion of PI(4,5)P_2_ by the PI(4,5)P_2_ phosphatase Ci-VSP released tubbyCT from the clusters, similar to dissociation of the domain from the bulk membrane (**Fig. 5A**), providing direct evidence for an essential role of PI(4,5)P_2_ in localizing tubbyCT to ER-PM junctions. Relative (fractional) decrease in tubbyCT residence in the junctions upon activation of the phosphatase was slightly but significantly less compared to the bulk membrane (**Fig. 5B**), which is consistent with the domain’s affinity to junctional E-Syt3. PI(4,5)P_2_-related effects may also play a role, in particular a higher local PI(4,5)P_2_ concentration or impaired access of Ci-VSP to its substrate in the ER-PM junction. Reduction in PI(4,5)P_2_ content in the PM by a pharmacological approach had a similar effect. PI(4,5)P_2_ is generated at the PM by sequential phosphorylation of PI to PI(4)P, mainly catalyzed by PI4KIIIα, and subsequently to PI(4,5)P_2_ by PI4P5 kinases (reviewed, e.g. in (Pemberton et al., 2019)). We inhibited this pathway by incubation with well-characterized PI4KA kinase inhibitors. Incubation with either the non-selective PI4K inhibitor phenylarsine oxide (PAO) or the highly selective PI4KIIIα inhibitor GSK-A1 (Bojjireddy et al., 2014) resulted in diminished association of tubbyCT to ER-PM junctions (**Fig. 5C-F**), corroborating the role of PI(4,5)P_2_ in recruitment of tubbyCT to these compartments.

**Figure 5.**
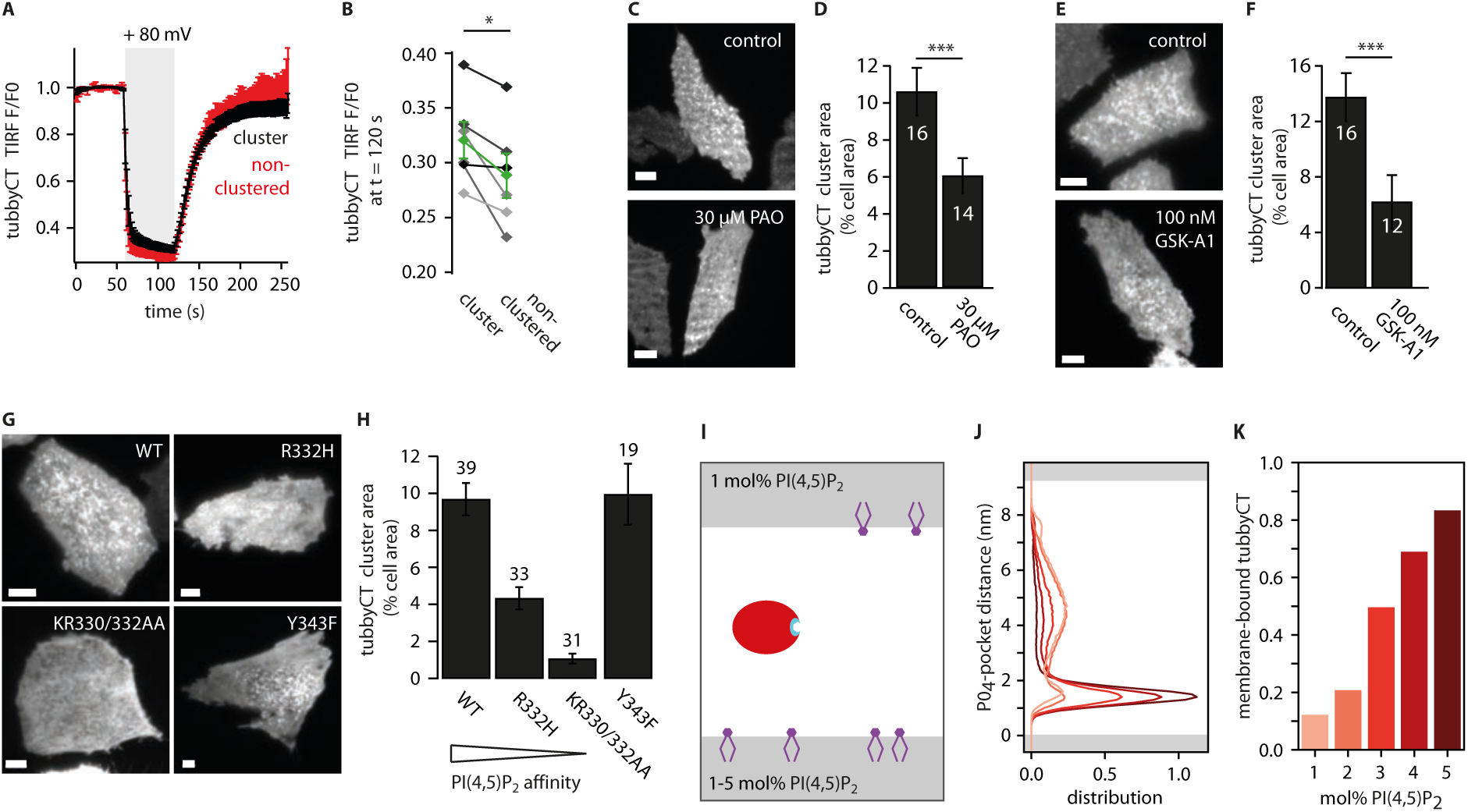
PI(4,5)P_2_ is essential for recruitment of tubbyCT into ER-PM junctions. **(A)** GFP-tubbyCT membrane dissociation dynamics in clustered versus non-clustered regions induced by PI(4,5)P_2_ depletion by activation of co-expressed Ci-VSP. Fluorescence amplitude obtained from TIRF imaging of whole-cell voltage-clamped cells (mean ± SEM; n = 6 CHO cells). **(B)** Comparison of residual tubbyCT membrane association after maximal Ci-VSP activation in the clustered and non-clustered regions, shown for the same individual cells analyzed in (A) with mean (± SEM) shown in green (paired t test, p = 0.0275). **(C)** Representative TIRF images of GFP-tubbyCT membrane distribution in CHO cells under control condition and after 30 min incubation with PAO (30 µM). Scale bars, 5 µm. **(D)** Average cell surface area occupied by tubbyCT clusters (mean ± SEM; p = 0.00695). **(E)** Representative TIRF images of GFP-tubbyCT membrane distribution in control CHO cells and after 10 min incubation with GSK-A1 (100 nM). **(F)** Average cell surface area occupied by tubbyCT clusters (mean ± SEM; p = 0.00578). **(G)** Representative TIRF images of CHO cells expressing GFP-tubbyCT wild-type (WT) and mutants R332H, KR330/332AA, and Y343F. **(H)** Cell area occupied by GFP-tubbyCT clusters analyzed from images as in (G). **(I)** Molecular dynamics simulations of tubbyCT binding to PI(4,5)P_2_ at the coarse-grained Martini level. Schematic simulation setup using a POPC bilayer (grey) doped with different fractions of PI(4,5)P_2_ (purple) in each leaflet. TubbyCT (red) was initially placed in the water phase between both leaflets. To analyse the binding affinity, the distance between the PI(4,5)P_2_ binding pocket (cyan) and the PO_4_ plane of the leaflet was analysed. **(J)** Histogram of the PO_4_-pocket distance using different concentrations of PI(4,5)P_2_ in the lower leaflet. The PI(4,5)P_2_ concentrations were ranging from 1-5 mol% (from light to dark red). **(K)** Membrane-bound population of tubbyCT localized at the lower leaflet containing an increasing PI(4,5)P_2_ concentration. In case of the setup with 1 mol% PI(4,5)P_2_, the population average of both leaflets (which both contained 1 mol% of PI(4,5)P_2_) was employed.

Further, we analyzed the junctional residency of tubbyCT mutants with reduced PI(4,5)P_2_ affinity. The semi-conservative mutation R332H affects one of the positive charges in the PI(4,5)P_2_ binding pocket (Santagata et al., 2001), which results in reduced PI(4,5)P_2_ affinity evidenced by poor membrane association (**Suppl. Fig. S4**; also see (Leitner et al., 2019; Quinn et al., 2008)). As shown in **Fig. 5G**, mutation R332H strongly reduced localization of tubbyCT to clusters (i.e. ER-PM junctions). Further reduction of PI(4,5)P_2_ affinity by the double mutation KR330/332AA (**Suppl. Fig. S4**), almost entirely abolished clustering (**Fig. 5G,H**). In contrast, a control mutation located within the lipid binding pocket but without substantially affecting PI(4,5)P_2_ affinity (**Suppl Fig. S4**; (Santagata et al., 2001)) did also not impact the clustering behavior (**Fig. 5G,H**).

While all of these findings are consistent with recruitment of tubbyCT into the ER-PM junctions by a local PI(4,5)P_2_ accumulation, they do not immediately explain why this would be specific to this particular PI(4,5)P_2_-binding domain. In order to understand how the interaction with PI(4,5)P_2_ might drive tubbyCT into clusters we took a closer look at its binding to PI(4,5)P_2_ by performing coarse-grained molecular dynamics simulations. Specifically, we simulated binding of tubbyCT to membrane domains containing different PI(4,5)P_2_ concentrations (**Fig.5I**) that compete for tubbyCT binding. **Fig5J** presents the observed distribution of membrane proximity of the major PI(4,5)P_2_ binding pocket of tubbyCT to two membrane compartments doped with different PI(4,5)P_2_ concentrations. Notably, a moderately higher concentration (2-5 mol%) resulted in the supralinear accumulation of tubbyCT at this membrane relative to the second compartment containing a lower (1 mol%), but physiologically plausible concentration (McLaughlin and Murray, 2005) of PI(4,5)P_2_ (**Fig. 5K**).

Of note, these simulations revealed a second, previously unrecognized but equally important PI(4,5)P_2_ interaction site at tubbyCT’s membrane interaction surface (Thallmair et al., 2020), which may explain this highly non-linear PI(4,5)P_2_-binding behavior not observed with PLCδ1-PH. Thus, moderately higher PI(4,5)P_2_ concentrations in the ER-PM junctional domain could strongly attract tubbyCT, but not other PI(4,5)P_2_-binding domains such as PLCδ1-PH or ENTH. In conclusion, a coincidence detection mechanism involving interaction with both E-Syt3 and PI(4,5)P_2_ mediates localization of tubbyCT to ER-PM junctions.

### TubbyCT recruitment reveals local synthesis of PI(4,5)P_2_ at ER-PM junctions during PLC activation

Given the dependence of tubbyCT localization on both, E-Syt3 and PI(4,5)P_2_, the pronounced additional recruitment into ER-PM junctions during Gq/PLC activity could be brought about by increases either in local E-Syt3 abundance or in local PI(4,5)P_2_ concentration.

It is known that PLC-induced increase in cytosolic Ca^2+^ recruits E-Syt1 to ER-PM junctions (Chang et al., 2013; Giordano et al., 2013; Idevall-Hagren et al., 2015), which was also observed in our experimental model, as shown in **Fig. 6A**. In contrast, E-Syt-3 did not accumulate substantially at the membrane in response to stimulation of the Gq-coupled M1 receptor. We observed only a minor and delayed increase in E-Syt3 fluorescence. Because this time course differs from the behavior of tubbyCT under the same conditions both in terms of amplitude and kinetics (**Fig. 6A**), E-Syt3 dynamics are unlikely to drive the recruitment of tubbyCT.

**Figure 6.**
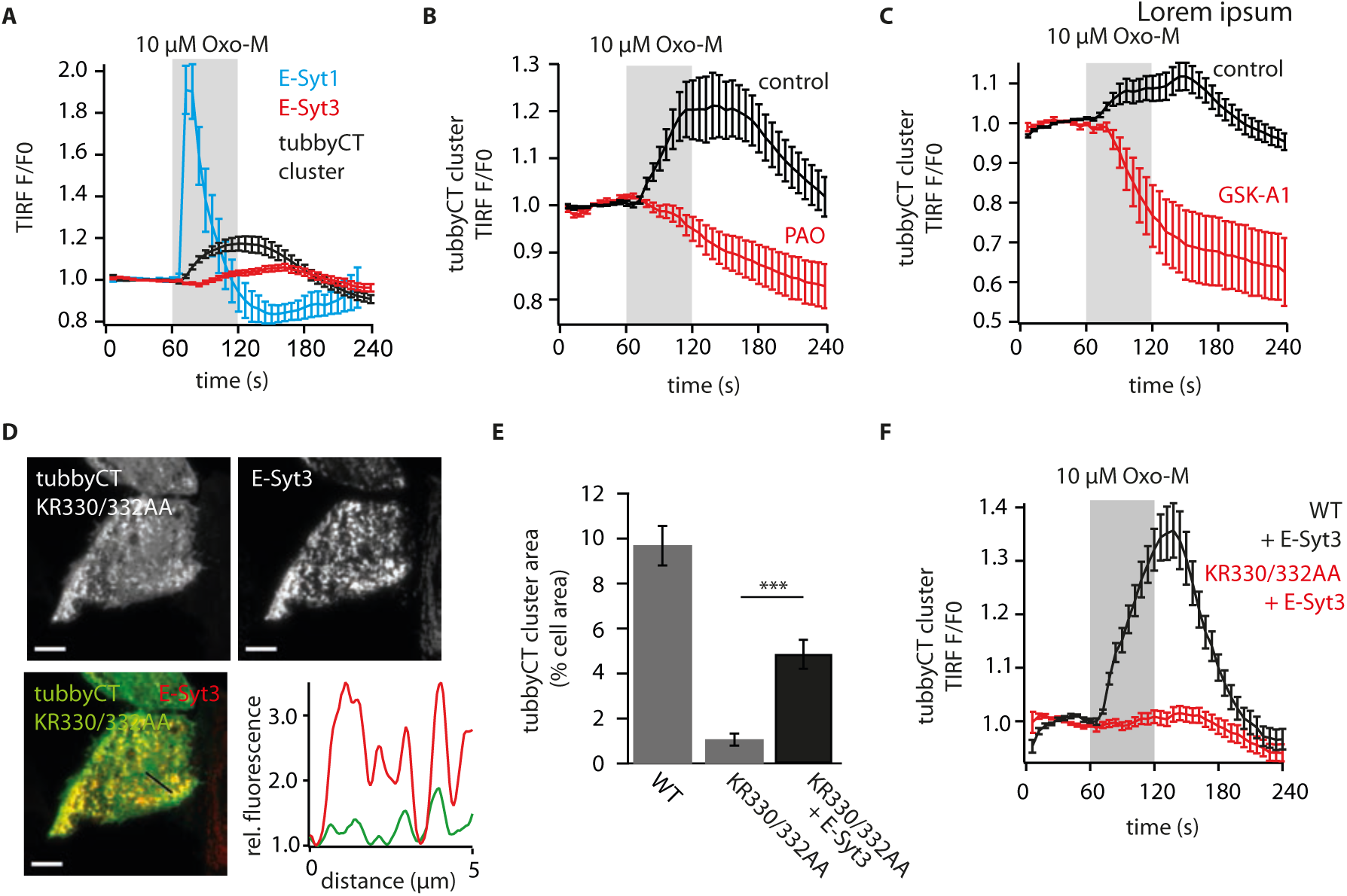
Spatial PI(4,5)P_2_ pool dynamics require local resynthesis at ER-PM junctions. **(A)** Membrane association dynamics of RFP-E-Syt1, RFP-E-Syt3 and GFP-tubbyCT in CHO cells in response to activation of PLC-β by stimulation of co-expressed M1R (mean ± SEM from, 19, 47, and 49 cells, respectively). **(B, C)** Effect of PI4K inhibitors on recruitment of tubbyCT into ER-PM junctions in response to PLC-β signaling. CHO cells expressing GFP-tubbyCT were pre-incubated with PAO (30 µM for 30 min, B) or GSK-A1 (100 nM for 10 min, C), and stimulation of M1R was done in the continued presence of the compounds. TubbyCT fluorescence (mean ± SEM) in cluster regions identified in resting state was recorded by TIRF-M and is shown normalized to signal strength before stimulation. (B: control, n = 40 cells; PAO: n = 24; C: control, n = 39; GSK-A1, n = 25) **(D)** Localization of the PI(4,5)P_2_-insensitive mutant of GFP-tubbyCT to ER-PM junctions induced by overexpression of E-Syt3. Representative TIRF images of CHO cells co-expressing GFP-tubbyCT K330AR332A and RFP-E-Syt3. Scale bar = 5 µm. Line profiles were derived from track highlighted in merged image and are plotted with fluorescence intensities normalized to minimal values (GFP-tubbyCT KR330/332AA, green; RFP-E-Syt3, red). **(E)** Quantitative analysis of cluster (ER-PM) localization of tubbyCT-K330AR332A E-Sy3 Cell area occupied by tubbyCT wild-type and K330AR332A clusters. Grey bars are replotted from Figure 5H. (KR330/332AA + RFP-E-Syt3: n = 43 CHO cells; Student’s t test: p = 0.00000122). **(F)** Dynamic recruitment of GFP-tubbyCT (WT, black) and GFP-tubbyCT KR330/332AA (red) into clusters in response to activation of PLC-β. CHO cells were co-transfected with tubbyCT constructs and RFP-ESyt3, and the TIRF signal was analyzed in detected clusters only (mean ± SEM; WT, n = 14; KR330/332AA, n = 34).

Instead, PI(4,5)P_2_ dynamics may cause the stimulus-induced recruitment of tubbyCT into the ER-PM-junctions. To test this idea, we monitored tubbyCT behavior in response to receptor stimulation while manipulating PI(4,5)P_2_ synthesis. Since it is established that during strong PLC activation resynthesis of PI(4,5)P_2_ requires the synthesis of PI(4)P in the PM (Bojjireddy et al., 2014), we again used inhibitors of PI4K to block any PI(4,5)P_2_ supply while activating the Gq/PLC coupled M1 receptor. In the presence of PAO, recruitment of tubbyCT was abolished and the domain instead dissociated from the ER-PM junctions (**Fig. 6B**). Similarly, GSK-A1 (Bojjireddy et al., 2014) abrogated recruitment of tubbyCT, ultimately resulting in full release of the domain from the ER-PM junctions upon receptor activation (**Fig. 6C**).

A tubbyCT mutant with strongly reduced PI(4,5)P_2_ affinity (KR330/332AA), while lacking clustering under basal conditions (cf., Fig. 5), could be recruited to some degree into ER-PM junctions by over-expression of E-Syt3 (**Fig. 6D,E**) but not by E-Syt1 or E-Syt2 (**Suppl. Fig. 5**), reconfirming the specific interaction with E-Syt3. However, in contrast to wildtype tubbyCT, we observed no further recruitment of the mutant upon receptor-mediated activation of PLCβ (**Fig. 6F**). Therefore, direct PI(4,5)P_2_ binding rather than E-Syt3 binding determines the dynamic behavior of tubbyCT in the ER-PM junctions.

Together, these results demonstrate that PI(4,5)P_2_ is the critical factor mediating PLC-dependent recruitment into ER-PM junctions. Specifically, our results show that in these domains the PI(4,5)P_2_ concentration increases during PI(4,5)P_2_ consumption by PLC, opposite to the PI(4,5)P_2_ dynamics in the bulk PM.

### PI(4,5)P_2_ production supports integrity of ER-PM junctions during Gq/PLCβ signaling

What might be the functional relevance of stimulated PI(4,5)P_2_ increase in ER-PM junctions? Among the junction-resident proteins, several, including the E-Syts, are tethered to the PM by binding to PI(4,5)P_2_. We therefore were interested in the dynamic membrane association of the E-Syts during PLCβ signaling. As described above (see **Fig. 6A**), E-Syt1 was strongly but transiently recruited to the junctions, which is known to be predominantly mediated by Ca^2+^ dynamics (Idevall-Hagren et al., 2015). In contrast, E-Syt2 and E-Syt3, lacking the Ca^2+^-binding C2 domain unique to E-Syt1 did not show pronounced PM recruitment when the Gq-coupled M1 receptor was stimulated. Instead, RFP-E-Syt2 showed a moderate and quickly reversible loss of membrane association and E-Syt3 PM association was only slightly affected by PLC signaling (**Fig. 7B** and **C**, respectively). When PI(4,5)P_2_ increase was prevented by blocking resynthesis with GSK-A1, membrane dynamics of all three isoforms were strongly altered (**Fig. 7A-C**). Thus, E-Syt1 recruitment was largely suppressed, and both E-Syt2 an E-Syt3 displayed pronounced loss of membrane association. Since attachment to the PM is mediated by binding of the terminal C2 domain to PI(4,5)P_2_ for all three isoforms (Giordano et al., 2013; Idevall-Hagren et al., 2015), this behavior is consistent with the pronounced loss of PI(4,5)P_2_ from the PM (cf. **Fig 6C**). Given the stable N-terminal insertion of E-Syts in the ER membrane, declining fluorescence at the PM as measured by TIRF-M indicated an increased distance between ER and PM and/or a loss of E-Syts from the junctions by lateral diffusion. In either case, blocking the local PI(4,5)P_2_ signal impacted on the integrity of ER-PM junctions, which in turn predicts impairment of homeostatic lipid transfer activity at these sites. Thus, the increased PI(4,5)P_2_ formation in the junctional PM compartment is essential to maintain or even enhance the functional state of E-Syts during PLCβ signaling.

**Figure 7.**
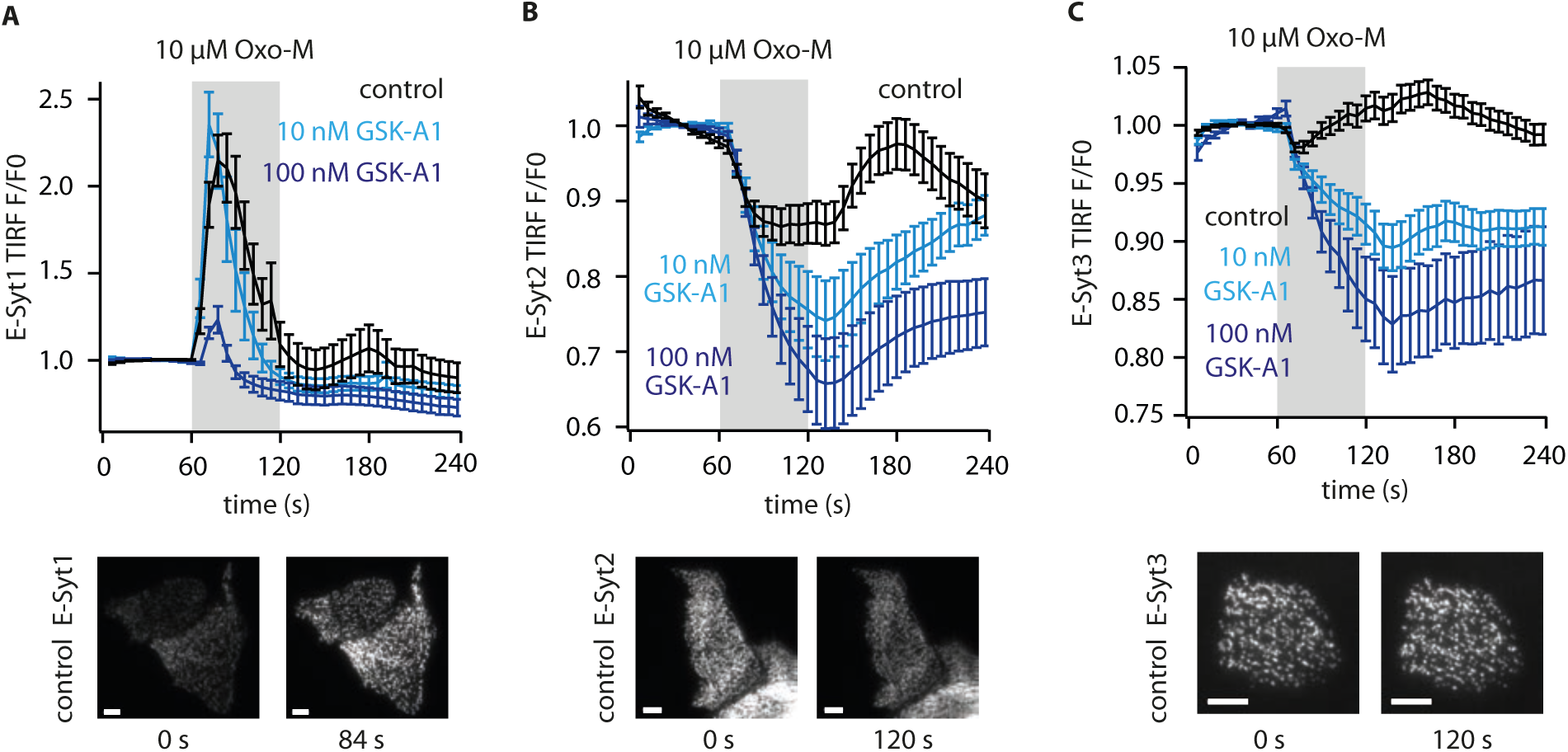
Local PI(4,5)P_2_ increase maintains membrane tethering by E-Syts during PLCβ signaling. **(A)** Influence of PI4K inhibitor GSK-A1 on E-Syt1 membrane association dynamics in response to activation of PLC-β. CHO cells expressing GFP-E-Syt1 were incubated in GSK-A1 (10 min) and subsequently imaged by TIRF microscopy during application of M1R agonist Oxo-M in the continued presence of GSK-A1. Representative images from control experiments without GSK-A1 before PLC-β activation (t = 0 s) and at maximal PLC-β response are shown in lower panels. Scale bar = 5 µm. (control, n = 20 cells; 10 nM GSK-A1, n = 16; 100 nM GSK-A1, n = 15). **(B)** Impact of GSK-A1 on E-Syt2 dynamics measured as described in (A). (control, n = 23; 10 nM GSK-A1, n = 16; 100 nM GSK-A, n = 16) **(C)** Impact of GSK-A1 on E-Syt3 dynamics measured as described in (A). (control, n = 42; 10 nM GSK-A1, n = 27; 100 nM GSK-A1, n = 26)

### Behavior of full-length tubby and TULP protein homologs

So far, we used the isolated tubby domain to uncover lipid dynamics at the ER-PM junctions. We wondered if association with ER-PM contact sites is also retained with the full-length tubby protein and whether this property is a general feature of members of the tubby protein family.

As shown in **Fig. 8A,B**, full-length tubby showed the same clustered distribution at the PM as the isolated tubbyCT domain, indicating that the native protein also preferably localizes to ER-PM junctions.

**Figure 8.**
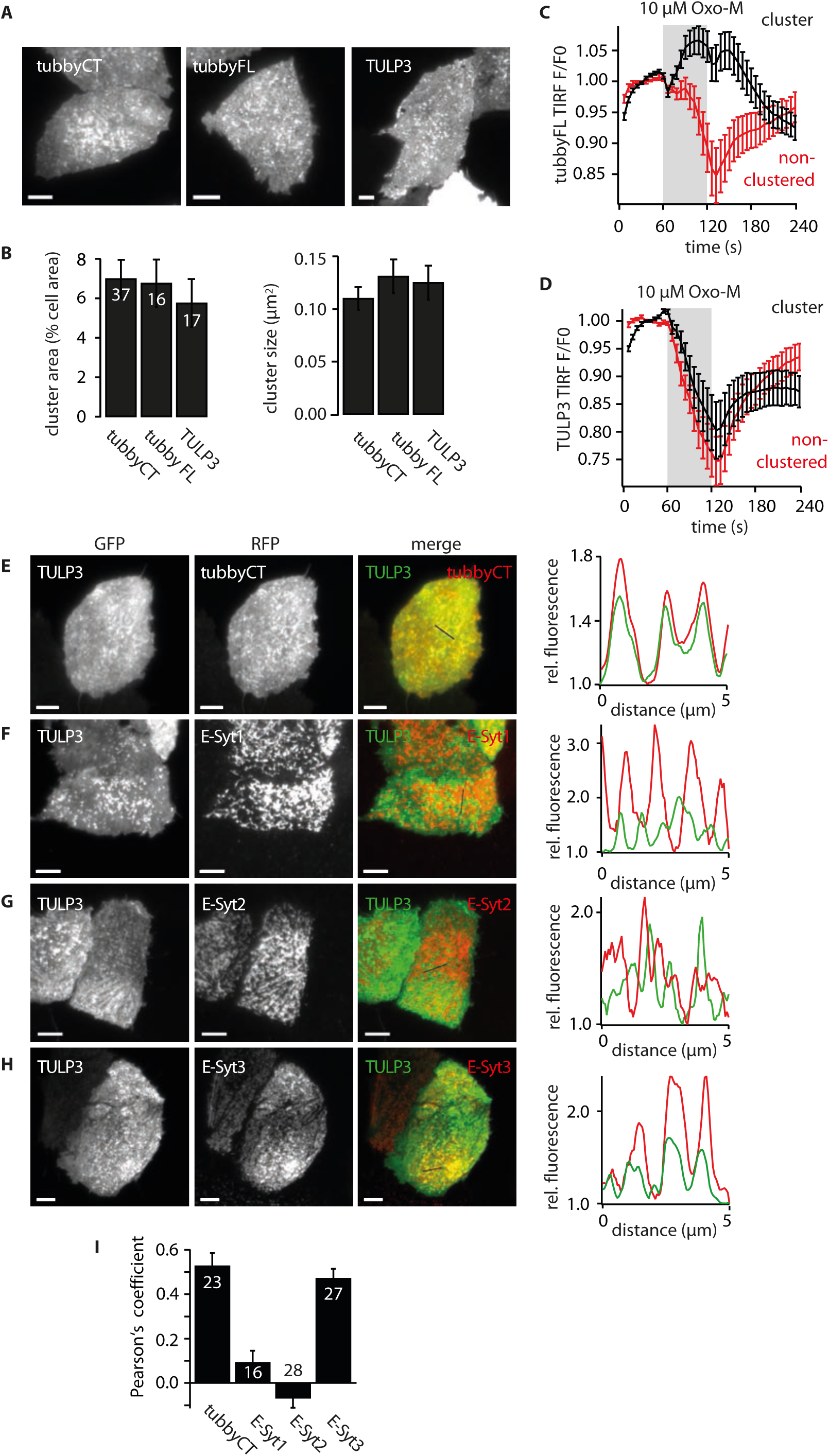
Localization of TULP proteins to ER-PM junctions. **(A)** Representative TIRF images of CHO cells expressing GFP-tubbyCT, full-length GFP-tubby (tubbyFL) or GFP-TULP3. Scale bar = 5µm. **(B)** Mean (± SEM) cluster area (left panel) and cluster size (right) from cells expressing the constructs shown in (A). **(C)** Membrane association dynamics of GFP-tubbyFL in response to activation of PLC-β by stimulation of co-expressed M1R, analysed separately for clusters (black) and non-clustered membrane regions (red; mean ± SEM from n = 15 cells). **(D)** Membrane association dynamics of GFP-TULP3 recorded as described in (C) (n = 17 cells). **(E)** Representative TIRF image demonstrating that TULP3 occupies the same domains enriched in tubbyCT. Right panel, fluorescence intensity line profiles (normalize to minimal intensities)for representative region indicated in merged fluorescence images. **(F-H)** Co-localization analysis of GFP-TULP3 with RFP-E-Syt1, RFP-E-Syt2, or RFP-E-Syt3, respectively. CHO cells co-expressing RFP-fused E-Syt isoforms with GFP-TULP3 were imaged using TIRF microscopy. Location of fluorescence intensity line profiles plotted in right panels is shown in merged fluorescence images. **(I)** Quantitative analysis of colocalization of TULP3 with tubbyCT and E-Syt isoforms using Pearson’s coefficient (mean ± SEM).

Tubby is the founding member of the family of tubby-like proteins (TULPs). Notably, TULP1-3 share the conserved C-terminal tubby domain (Mukhopadhyay and Jackson, 2011; Wang et al., 2018). We examined localization of TULP3, which has the most ubiquitous distribution across tissues in mammals (Mukhopadhyay and Jackson, 2011). TIRF imaging showed that in CHO cells, TULP3 has substantial PM localization which is organized in clusters highly similar to the distribution of tubby, both with respect to size and membrane area covered by the clusters **(Fig. 8A,B)**. However, unlike full-length tubby, which recapitulated the recruitment into junctions observed with its isolated C-terminal domain (**Fig. 8C**), TULP3 dissociated from the junctions upon activation of Gq/PLC signaling (**Fig. 8D**). Nevertheless, co-localization experiments with tubbyCT and with E-Syt isoforms (**Fig. 8E-I**) showed that TULP3 preferentially accumulates at E-Syt3-positive ER-PM junctions populated by tubby. Thus TULP3 may quantitatively differ in its affinity to PI(4,5)P_2_ or E-Syt3.

In conclusion, PI(4,5)P_2_-dependent association with ER-PM junctions appears to be a general feature of TULPs, intriguingly suggesting they also serve a yet to be discovered function at these central cellular hubs of lipid and Ca^2+^ metabolism.

## DISCUSSION

### An ER-PM junctional pool of PI(4,5)P_2_

Unexpectedly, we discovered that the tubby domain, by virtue of a coincidence detection mechanism involving binding to both PI(4,5)P_2_ and the tether protein E-Syt3 preferentially segregates into ER-PM junctions that contain E-Syt3. Since this local enrichment was dependent on PI(4,5)P_2_, it provided the first opportunity to selectively examine a junctional PI(4,5)P_2_ pool and its dynamics. Specifically, we have observed the seemingly paradoxical increase of PI(4,5)P_2_ concentration in the PM of these domains during PLCß signaling, when consumption of PI(4,5)P_2_ in the bulk PM is reported by various PI(4,5)P_2_ sensors.

This inverted PI(4,5)P_2_ dynamics in tubbyCT-positive ER-PM junctions as opposed to the bulk PM provides direct evidence for the spatial organization of two distinct pools of PI(4,5)P_2_ in the PM. Moreover, the local clustering of tubbyCT in resting cells indicates enrichment of PI(4,5)P_2_ also under basal conditions. While binding to ER-PM junction-resident E-Syt3 provides a straightforward explanation for basal tubbyCT enrichment, we observed pronounced reverse recruitment of truncated E-Syt3 lacking an ER-anchor into the junctions by tubbyCT suggesting a local enrichment of PI(4,5)P_2_ that is sufficient for concentrating tubbyCT and its interaction partner. Preferential recruitment of tubbyCT by PI(4,5)P_2_, which was observed only very subtly with PLCδ1-PH, can be attributed to its lower affinity compared to PLCδ1-PH, and tubbyCT’s cooperative binding of PI(4,5)P_2_ via two distinct binding sites located at the domain-membrane interface (Thallmair et al., 2020).

The question as to whether PM phosphoinositides are organized into distinct pools has attracted attention since long, as it may explain how a single molecule can instruct a multitude of different cellular processes with specificity (reviewed in (Doughman et al., 2003; Hammond, 2016)). Experimental evidence includes both functional assays as well as fluorescence imaging approaches to address the spatial organization of PI(4,5)P_2_ (and other PIs). As an experimentally particularly well-supported example, PI(4,5)P_2_ synthesis from PI(4)P at the PM is largely independent from bulk PI(4)P, indicating a separate functional pool of PI(4)P designated for the synthesis of PI(4,5)P_2_ (Bojjireddy et al., 2014; Hammond et al., 2012; Hammond et al., 2009; Nakatsu et al., 2012). Structurally, high resolution light microscopy and EM using PI(4,5)P_2_ -specific probes (either antibodies or PLCδ1-PH) have demonstrated heterogeneity and even labeling of well-defined PI clusters (Fujita et al., 2009; van den Bogaart et al., 2011; Wang and Richards, 2012). However, these approaches generally used fixation protocols that potentially affect the lateral distribution of PIs (Hammond, 2016; van Rheenen et al., 2005), and other careful studies using live-cell approaches failed to demonstrate substantial spatial heterogeneity (Ji et al., 2015; van Rheenen et al., 2005).

Enrichment of PI(4,5)P_2_ at ER-PM junctions has been postulated before (Maléth et al., 2014), however, direct evidence has not been provided so far (Chang and Liou, 2016).

### Local synthesis of the ER-PM junctional PI(4,5)P_2_ pool

How might the junctional buildup of PI(4,5)P_2_ arise? As shown by pharmacological inhibition of PI4K, the PI(4,5)P_2_ increase in ER-PM junctions during receptor activation requires de-novo synthesis from PtdIns. Because the PI(4,5)P_2_ concentration in the rest of the membrane drops simultaneously, the localized increase must arise from local synthesis in the PM of the junctions. This conclusion implies that the enzymatic equipment for both phosphorylation steps, i.e. both PI4K and PI4P5K, are localized to these junctions. PI4KIIIα, which is the major source of PI(4)P at the PM (Balla et al., 2007; Balla et al., 2005; Nakatsu et al., 2012) is assembled into an evolutionary conserved multi-protein complex that mediates PM targeting (Chung et al., 2015a; Lees et al., 2017b; Nakatsu et al., 2012). Both PI4P5Kα and PI4KIIIα can be assembled into an enzymatic complex by the scaffolding protein IQGAP1 (Choi et al., 2016). These interactions are consistent with the idea that both enzymes may also be targeted into ER-PM junctions by protein-protein interactions. Interestingly, in yeast, the homologous PI4K (Stt4) complex clusters in microscopic ’phosphoinositide kinase patches’ at the PM (Baird et al., 2008). However, despite a striking similarity of this cortical pattern in yeast with the distribution of ER-PM junctions (Nakatsu et al., 2012; Stefan et al., 2011), to our knowledge a correspondence of PI4K patches and ER-PM contact sites has not been shown.

In considering the signal that triggers the localized PI(4,5)P_2_ synthesis, we argue that sensing of global PI(4,5)P_2_ by a rheostatic PI(4,5)P_2_ sensor is unlikely to play an important role, since global depletion of PI(4,5)P_2_ via an alternative enzymatic pathway, i.e. by the voltage-dependent phosphatase (VSP) did not induce any detectable local increase of PI(4,5)P_2_ at the junctions (**Fig. 5A**).

Instead, we consider two possible mechanisms. First, the Gq/PLC pathway may provide a feedback signal that activates one or both of the kinases that generate PI(4,5)P_2_ from its precursor PtdIns. Obvious candidate signals may be the concomitant Ca^2+^ surge, which has been shown to affect ER-PM junctions by directly mediating recruitment of E-Syt1, or more indirectly PKC (Tóth et al., 2016) or the neuronal Ca^2+^-binding protein (NCS-1) previously suggested to activate PI4K in neurons (Delmas et al., 2005; Winks et al., 2005).

Alternatively, the local synthesis may simply be fed by the stimulated supply of its substrate, PtdIns, which is provided by transport from the ER by the PI transport proteins Nir2/Nir3 (Chang et al., 2013; Kim et al., 2013; Kim et al., 2015) and/or TMEM24 dynamically recruited into ER-PM junctions during signaling. This transport activity was already demonstrated to be fundamental for the resynthesis of the PM pools of PI(4,5)P_2_ (Chang and Liou, 2015; Kim et al., 2015). Given that such PtdIns transfer occurs primarily at the ER contact sites, it will provide an inherently local supply of substrate. This model of spatial metabolic channeling fits exceptionally well to the recent finding that PtdIns levels are actually very low in the PM(Pemberton et al., 2020; Zewe et al., 2020). Combined with that observation, our results thus provide evidence that PtdIns arriving from the ER is immediately turned over to PI(4)P and PI(4,5)P_2_ within the ER-PM contact zone, which may subsequently leave this domain by diffusion to restore the PM pools of phosphoinositides.

We note that this model of metabolic channeling can also account for a conundrum concerning phosphoinositide pool identity at the PM: Hammond and colleagues found that although it is clear that PI(4,5)P_2_ is synthesized from PI(4)P in the PM, depletion of bulk PI(4)P had surprisingly little impact on the PI(4,5)P_2_ concentration and on the recovery of PI(4,5)P_2_ following depletion by PLC (Hammond et al., 2012; Hammond et al., 2009). Thus, bulk PI(4)P appears to serve as a determinant of PM identity rather than as the substrate pool for PI(4,5)P_2_ synthesis (Hammond et al., 2012), while a distinct hidden pool of PI(4)P must be postulated as the precursor for PI(4,5)P_2_. If conversion of PI(4)P to PI(4,5)P_2_ primarily occurs at ER-PM junctions as suggested by our findings, the newborn PI(4)P in these domains may represent the cryptic pool serving PI(4,5)P_2_ resynthesis during PLC signaling.

### Functional relevance of local PI(4,5)P_2_ increase

Our data suggest that beyond channeling PI(4,5)P_2_ synthesis for sustained PLC-coupled receptor signaling, the local synthesis and increase of PI(4,5)P_2_ serves an important role in ER-PM junction functionality. The majority of the tethering proteins that establish or tighten the contact sites during signaling are ER-resident proteins but use PI(4,5)P_2_ and/or PI(4)P for anchorage to the PM, including the E-Syts (Giordano et al., 2013), ORPs (Chung et al., 2015b), TMEM24 (Lees et al., 2017a), and GRAMD proteins (Besprozvannaya et al., 2018). Further, recruitment of STIM1 in ER-PM junctions to trigger SOCE requires PI(4,5)P_2_ in the PM (Walsh et al., 2009). Therefore quite paradoxically, while the homeostatic ER-PM function should be most active during PLC signaling, the associated PI(4,5)P_2_ consumption would disrupt ER-PM tethering, homeostatic lipid transport activity, and SOCE. Thus, the local surge in PI(4,5)P_2_ serves to maintain and possibly even enhance the pivotal transport activities at ER-PM junctions. In fact, when we blocked the junctional PI(4,5)P_2_ increase, we observed pronounced loss of E-Syts from the junctions during signaling, indicating inactivation of the junctional lipid transfer.

As a word of caution it should be emphasized that the coincidence detection properties of the tubbyCT sensor limits our observations to ER-PM junctions populated by E-Syt3. The finding that tubbyCT does not localize to junctions earmarked by (overexpressed) E-Syt1 or 2, the diversity of tethering proteins identified so far, as well as direct evidence (e.g. (Besprozvannaya et al., 2018; Giordano et al., 2013)) indicate substantial heterogeneity of ER-PM junctions. Hence, we currently do not know if the junctional PI(4,5)P_2_ surge is a general feature of ER-PM junctions or whether it is restricted to those characterized by E-Syt3. In fact, the transient nature of E-Syt1 recruitment during onset of PLC signaling has been understood to reflect the loss of PI(4,5)P_2_ from the junctional PM (Idevall-Hagren et al., 2015; Saheki et al., 2016). However, our results show that inhibition of PI kinases further curtails degree and duration of recruitment of E-Syt1, indicating a similar local resynthesis of PI(4,5)P_2_ in junctions populated by E-Syt1.

### tubbyCT as a sensor for junctional PI(4,5)P_2_ dynamics

Taken together, the specific coincidence detection properties of tubbyCT allow monitoring the local PI(4,5)P_2_ pool and its dynamics during signaling specifically for those ER-PM junctions enriched in E-Syt3. However, the very same coincidence detection principle may be exploited to address PI dynamics in different types of membrane contact sites. To this end, we note the obvious potential to generate versatile, either general, or subtype-specific sensors for junctional phosphoinositides by fusing junctional targeting motifs to the various specific PI sensors that previously have been exploited for the examination of global PI dynamics. Indeed, engineered coincidence detectors for phosphoinositides revealed local PI dynamics at endocytic structures (He et al., 2017). There, a clathrin-binding module was fused to various PI-selective domains including to the PI(4,5)P_2_ -specific PH domain of PLCδ1, and subsequently used successfully to detect transient accumulation of PI(4,5)P_2_ during endocytosis events.

### Complexity and problems of commonly used PI(4,5)P_2_ biosensors

It was noted previously that tubbyCT is reluctant to dissociate from the PM during PLC activity (e.g., (Szentpetery et al., 2009)), and this had occasionally been attributed to a high PI(4,5)P_2_ affinity. However, quantitative titration of PI(4,5)P_2_ by a voltage-sensitive PI(4,5)P_2_ phosphatase (VSP) demonstrated unequivocally that its PI(4,5)P_2_ affinity is actually lower compared to the popular sensor domain, PLCδ1-PH (Halaszovich et al., 2009; Leitner et al., 2019). Molecular dynamics simulations support this lower affinity (Thallmair et al., 2020). The recruitment of tubbyCT into ER-PM junctions mediated by interaction with E-Syt3 and PI(4,5)P_2_ elevation in these domains during PLC signaling now conclusively explains these seemingly contradictory data.

Moreover, we recently discovered that tubbyCT binds PI(4,5)P_2_ with two structurally distinct binding sites (Thallmair et al., 2020), which may further contribute to preferential residence in PI(4,5)P_2_-enriched ER-PM junctions. Thus, together with the co-detection of E-Syt3, tubbyCT displays complex coincidence detection properties that need to be taken into account when interpreting its membrane association dynamics in terms of PI(4,5)P_2_ concentration changes. Notwithstanding, we have recently succeeded in using tubbyCT as a probe for neuronal PI(4,5)P_2_ dynamics in near-native hippocampal brain slice preparations. Different from its behavior in certain cell lines, tubbyCT readily dissociated from the PM following activation of muscarinic and glutamatergic Gq-coupled receptors, revealing neurotransmitter-dependent PI(4,5)P_2_ dynamics (Hackelberg and Oliver, 2018). Most likely, these neurons lack substantial expression of E-Syt3, which is consistent with RNAseq data indicating predominant E-Syt2 expression in the hippocampal formation of the mouse, but little E-Syt3 (Sjöstedt et al., 2020).

It should be emphasized that the use of the popular PI(4,5)P_2_ sensor, PLCδ1-PH, is severely compromised by similarly complex properties, namely the high affinity to IP3. Therefore, dissociation into the cytosol upon PLC signaling may result from the IP3 signal rather than depletion of PI(4,5)P_2_ (for a comprehensive discussion, see (Hammond and Balla, 2015)). Thus, although genetically encoded sensors have been widely used and have been instrumental in discovering many aspects of PI(4,5)P_2_ signaling and metabolism, quite surprisingly there is currently no well-characterized sensor that does not suffer from ambiguity in reporting PI(4,5)P_2_ concentrations. We recently characterized in some detail another structurally unrelated PI(4,5)P_2_ sensor domain, the ENTH domain from epsin. Here, membrane binding appears to depend on PI(4,5)P_2_ in a more straightforward manner (Leitner et al., 2019). However, it suffers from rather low membrane localization at rest, limiting the signal-to-noise ratio, particularly in confocal microscopy.

### Biological function of tubby (and TULP) localization

The preferred localization to ER-PM junctions was not only observed with the isolated tubbyCT domain, but was also prominent for the full-length tubby protein and for its homolog, TULP3. The conservation of this behavior in the TULP protein family, as well as the specificity implied by interaction with E-Syt3, suggest a specific role related to the function of ER-PM junctions. While such a mechanism remains to be determined, we point out that TULP proteins may either be effectors of PI(4,5)P_2_ dynamics in the ER-PM contact site, or vice versa they may have a role in controlling activity, including lipid metabolism, at these sites.

So far, TULP proteins have a well-established role in the selective transport of G-protein-coupled receptors (GPCRs), channels, enzymes, and signaling proteins into cilia (Badgandi et al., 2017; DiTirro et al., 2019; Han et al., 2019; Legué and Liem, 2019; Sun et al., 2012). Mistargeting of GPCRs and the ensuing ciliary dysfunction can explain at least some of the phenotypes observed in the tubby mouse mutant, which include obesity, retinal degeneration and hearing loss (reviewed in (Mukhopadhyay and Jackson, 2011)). Our new findings should stimulate the reconsideration of additional cellular mechanisms underlying the pathophysiology observed upon deletion of tubby family proteins. Considering a possible connection between ciliary function and ER-PM junctional localization, recruitment into ER-PM junctions may simply serve to sequester TULPs away from the cilium, thereby antagonizing and thus regulating ciliary signaling.

Intriguingly, a recent study showed that E-Syt3 is specifically expressed in hypothalamic nuclei involved in energy homeostasis and food intake (Zhang et al., 2020), and thus shows a striking overlap with neuronal tubby expression (including the arcuate nucleus)(Kleyn et al., 1996). Moreover, E-Syt3 impacts on diet-induced obesity, where increased E-Syt3 levels promoted obesity while loss of E-Syt3 antagonized obesity (Zhang et al., 2020). Given the cellular interaction discovered in this study, we speculate that both proteins may interact in the same pathway in regulating neuronal activity in energy homeostasis. In fact, both tubby and E-Syt were implicated in regulating POMC-derived transmitters in this neural circuitry (Guan et al., 1998; Zhang et al., 2020). Although entirely speculative at the moment, it is tempting to propose that E-Syt3 might sequester tubby in these neurons and thereby downregulate its function, explaining why the effects of increased E-Syt3 abundance resemble the consequences of loss of tubby.

In summary, our current findings emphasize the pivotal role that ER-PM junctions have in phosphoinositide homeostasis and signaling in the entire cell, and make a strong case for spatially compartmentalized PI signaling. In particular, they substantiate the concept of spatial metabolic channeling of PI synthesis in ER-PM junctions. Further, our observations suggest a novel, yet to be uncovered, function of tubby family proteins intimately linked to the biology of ER-PM contact sites.

## METHODS

### Cell culture

CHO dhFr^-^ cells were cultured in MEM Alpha medium (gibco, ThermoFisher Scientific, Waltham, US), COS-7 and MDCK cells in DMEM GlutaMAX™-I medium (gibco), both supplemented with 10% fetal calf serum, 1% penicillin and 1% streptomycin. Cells were kept at 37°C and 5 % CO2. They were seeded on glass bottom dishes (Figs. 1A-B, E-G, 3G-H, 5A-D, 6B) or in glass bottom µ-slide VI^0.5^ flow chambers (ibidi, Martinsried, Germany; Figs. 1C-D, G, Fig. 2, Fig. 3A-F, Fig. 4C-K, Fig. 5E-H, Fig. 6A, C-F, Fig. 7, Fig. 8 and Suppl. Figs. 1-5) for imaging experiments and on polystyrene dishes for protein extraction. Two days after seeding, cells were transfected using JetPEI® DNA transfection reagent (CHO dhFr^-^ cells, Polyplus Transfection, Illkirch-Graffenstaden, France) or lipofectamin®2000 reagent (COS-7 and MDCK cells, Invitrogen, Carlsbad, US). Experiments were performed 24h (TIRF imaging) or 48h (protein extraction) post-transfection. For all TIRF experiments involving the activation of PLCβ, cells were co-transfected with the respective constructs plus the human muscarinic receptor 1 (M1R).

### Molecular biology

Expression constructs used for transfection were: human M1R (NM_000738.2) in pSGHV0; human PLCδ1-PH (AA 1-170; NM_006225.3) in pEGFP-N1; rat epsin1-ENTH (AA 1-158) in pEGFP-N1 (NM_057136.1); Ci-VSP in pRFP-C1 (AB183035.1); mouse tubbyCT (AA 243-505) in pEGFP-C1 and pRFP-C1 (NM_021885.4); mouse tubbyFL in pEGFP-C1 (NM_021885.4); Lyn11-FRB (GCIKSKGKDSA) in pRFP-N1; human E-Syt1 in pEGFP-C1 (NM_015292; (Giordano et al., 2013)); human E-Syt2 in pEGFP-C1 (NM_020728.2, (Giordano et al., 2013)); human E-Syt3 in pEGFP-C1 (NM_031913.4, (Giordano et al., 2013); human Nir2-mcherry (Chang and Liou, 2015); human Nir3-cherry (AB385472, (Chang and Liou, 2015)); human TULP3 in pGLAP1 (NM_003324.4; (Badgandi et al., 2017)), pRFP-C1 and pEGFP-C1 as control vectors. RFP-tagged E-Syt1-3 constructs were obtained by subcloning E-Syt1-3 into pRFP-C3. CFP-tagged E-Syt3 construct was generated by fluorophore exchange of pEGFP-C1-E-Syt3 construct. Myc-tag was N-terminally added to tubbyCT via PCR amplification and sub-cloned into pcDNA3.1. Myc-PLCδ1-PH construct was generated by exchange of the tubbyCterm portion of pcDNA3.1-myc-tubbyCT with PLC1δ1-PH. N-terminally myc-tagged E-Syt1 was achieved via tag exchange of pEGFP-E-Syt1. In all myc-tagged constructs original start codon was removed by PCR. E-Syt1ΔHP (aa 125-1105), E-Syt2ΔHP (aa 106-846), E-Syt3δHP (aa 104-887), E-Syt3C2ABC (aa 296-887), E-Syt3C2BC (aa 439-887), E-Syt3C2C-L (aa 568-887), and E-Syt3C2C (aa 745-887) constructs were generated via PCR amplification from respective full-length constructs and subsequent sub-cloning into pRFP-C1 using BsrG1 and Sal1 (E-Syt1ΔHP), BglII and BamH1 (E-Syt2ΔHP) and HindIII and KpnI (E-Syt3 deletion constructs) restriction sites, respectively. pEGFP-C1-tubbyCT R332H and pEGFP-C1-tubbyCT Y343F were generated using QuikChange II XL Site-Directed mutagenesis kit (Stratagene, Agilent Technologies, Waldbronn, Germany). pEGFP-C1-tubbyCT KR330/332AA was generated by mutagenesis PCR using PfuUltra II Hotstart PCR Master Mix (Agilent Technologies, Santa Clara, US).

### Co-immunoprecipitation

CHO dhFr^-^ cells expressing myc- and GFP-fused constructs were washed in PBS and subsequently lysed in 50 mM Tris, 150 mM NaCl, 10 mM EDTA, 1% Triton X-100, pH = 7.2, 1% protease inhibitor cocktail (Roche) for 30 min. Cell debris was separated from lysate by centrifugation (21000 g; 4 °C; 20 min). 800 ng total protein was diluted in 600 µl dilution buffer (10 mM Tris, 150 mM NaCl, 0.5 mM EDTA, pH = 7.5) and used for immunoprecipitation (IP) by αGFP nanobodies covalently bound to agarose beads (GFP-Trap®_A, Chromotech, Planegg-Martinsried, Germany). 25 µl beads were washed three times in dilution buffer and then incubated with diluted cell lysates for 1 h at 4°C. After protein binding, beads were isolated by centrifugation (2500 g, 4 °C, 2 min) and 3x washed in dilution buffer followed by 3 washing steps in high Na+ buffer (10 mM Tris, 500 mM NaCl, 0.5 mM EDTA, pH = 7.5). After final centrifugation, immunocomplexes were detached from beads by incubation in 2x Lämmli buffer (20% v/v glycerol, 100 mM Tris, 130 mM DTT, 6.5% w/v SDS, 0.013% (w/v) bromphenol blue) (95 °C, 10 min). Supernatant was separated from beads by centrifugation (2500 g, 2 min) and used for Western Blot analysis.

Myc- and GFP-tagged proteins were detected by rabbit polyclonal IgG anti-GFP (FL) (1:200; sc-8334 Santa Cruz Biotechnology, Dallas, US) and mouse monoclonal IgG anti-c-Myc (1:200; sc-40 Santa Cruz Biotechnology) primary antibodies and goat αrabbit (IRDye® 800CW) (1:5000, 926-32211, Li-cor Biosciences, Bad Homburg, Germany) and donkey αmouse (IRDye® 800CW) (1:5000, 926-32212, Li-cor Biosciences) secondary antibodies, respectively. For each Co-IP experiment a negative control lysate of cells expressing myc-tubbyCT only was added. For calculation of relative Co-IP values the band intensity of this negative control was subtracted from α-myc band intensities and resulting Co-IP values were normalized to immunoprecipitated αGFP band intensities.

### TIRF microscopy

Experiments shown in Figs. 1A-B, 1E-G, 3G-H, 5A-D, and 6B were performed as previously described (Halaszovich et al., 2009). In brief, imaging was done on a BX51WI upright microscope (Olympus, Hamburg, Germany) provided with a TIRF condenser (Olympus) and a LUMPlanFI/IR 40×/0.8-numerical aperture water immersion objective (Olympus). Laser excitation occurred at 488 nm (Picarro, Sunnyvale, CA), emission light passed a 530 nm-long-pass emission filter and was detected by a CCD camera (TILL Photonics GmbH, Gräfelfing, Germany). Image acquisition was controlled by TILLvisION software (TILL Photonics GmbH). All other TIRF experiments were done on a Dmi8 upright microscope (Leica, Wetzlar, Germany) equipped with an Infinity TIRF module (Leica), a HC PL APO 100x/1.47 OIL objective (Leica) and a widefield laser (Leica). GFP-, RFP- and CFP-fluorescence was excited at 488 nm, 561 nm and 405 nm, respectively. Corresponding emission light passed GFP-T (505-555 nm), DS-Red-T (590-650 nm) and CFP-T (460-500 nm) emission filters (Leica). Images were acquired with an ORCA-Flash4.0 C13440-20C camera (Hamamatsu photonics, Hamamatsu, Japan) controlled by LAS X software (Leica).

During imaging cells were transfused with extracellular solution (5.8 mM KCl, 144 mM NaCl, 0.9 mM MgCl_2_, 1.3 mM CaCl_2_, 0.7 mM NaH_2_PO_4_, 5.6 mM D-glucose, 10 mM HEPES, pH = 7.4). In experiments combining TIRF-M and patch-clamping images were taken every 4 s and in all other time-resolved TIRF experiments images were taken every 6 s.

### Wide field fluorescence microscopy

Experiments were performed on a Dmi8 upright microscope (Leica, Wetzlar, Germany). Images were acquired with an ORCA-Flash4.0 C13440-20C camera (Hamamatsu photonics, Hamamatsu, Japan) controlled by LAS X software (Leica). For determination of membrane localisation of GFP-tubbyCT constructs, CHO cells were co-transfected with a catalytically inactive Ci-VSP (RFP-Ci-VSP C363S) as membrane marker. Line profiles across the cells were analysed. GFP fluorescence intensities at Ci-VSP C363S membrane peaks were averaged and normalized to average cytosolic fluorescence.

### Patch Clamp electrophysiology

Whole-cell patch clamp experiments for activation of Ci-VSP were performed as described previously (Halaszovich et al., 2009; Leitner et al., 2019). Briefly, voltage clamp recordings were done simultaneous to TIRF imaging with an EPC 10 amplifier controlled by PatchMaster software (HEKA Elektronik, Lambrecht, Germany). Patch pipettes were pulled from borosilicate glass (Sutter Instrument Company, Novato, CA, USA) and had an open pipette resistance of 2-3 MΩ after back-filling with intracellular solution containing (mM) 135 KCl, 2.41 CaCl_2_, 3.5 MgCl_2_, 5 HEPES, 5 EGTA, 2.5 Na_2_ATP, 0.1 Na_3_GTP, pH 7.3 (with KOH), 290-295 mOsm/kg. Series resistance (R_s_) typically was below 6 MΩ. Cells where held at -60 mV and stepped to +80 mV to activate phosphatase activity of Ci-VSP for 60 s.

### Chemicals

Oxotremorin-M (Oxo-M, Tocris Bioscience, Bristol, UK) was prepared in a 10mM stock solution in water and diluted to 10 µM in extracellular solution. The PI4 kinase inhibitors GSK-A1 (SYNkinase, Parkville, Australia) and phenylarsine oxide (PAO, Sigma Aldrich, St Louis, US) were dissolved in 1 mM (GSK-A1) and 100 mM (PAO) stock solutions in DMSO and diluted to their working conditions (10 nM and 100 nM GSK-A1; 30 mM PAO) in extracellular solution. Cells were incubated in PI4 kinase inhibitors for 10 min (GSK-A1) and 30 min (PAO) prior to imaging.

### Data analysis

For experiments shown in Fig. 1A-B, G and Fig. 6B image analysis was done with TILLvisION software (TILL Photonics GmbH) and IGOR Pro (WaveMetrics, Lake Oswego, OR, USA). For experiments shown in Fig. 1C,F,G, Figs. 2-3, Figs. 4D-K, Figs. 5, Figs. 6A, C-F, and Figs. 7-8 image analysis was done with ImageJ software. Pearson coefficients were calculated using Coloc2 plug-in. Sectioning algorithm for detection of clustered and non-clustered cell regions: back-ground subtracted images were median-filtered (r = 0.9 µm), subtracted from the back-ground subtracted original image and thresholded using RenyiEntropy threshold. Pre-cluster regions of interest (ROIs) were generated from resulting binary images. Pre-clusters were detected for each of the 40 images of the time series. Pixel assigned at least twice (TIRF microscope, 100x objective, Leica) or 6x (TIRF microscope, 40x objective, Olympus) to pre-clusters were added to the final cluster ROI. Pixels with a distance of 1.8 µm to those pixels detected at least twice as pre-clusters were added to the non-cluster ROI. Fluorescence dynamics in the clustered and non-clustered region was analysed. For analysis of cluster size and cluster coverage of the cell, only baseline clusters were taken into account.

Analysis and statistical evaluation of obtained imaging data was done with IGOR Pro (WaveMetrics, Lake Oswego, OR, USA). Data are displayed as mean ± SEM. For comparison of two groups Student’s t tests were performed. Values derived from the same cells were analysed using paired Student’s t tests. Single sample t testing (H0 = 0) was done for co-localization analysis of Pearsons’s coefficients. Asterisks indicate significance levels: *: p < 0.05, **: p < 0.01, ***: p < 0.001. Respective p values are also given in the figure legends.

### Coarse-grained molecular dynamics simulations

All simulations were performed using the open-beta version of the coarse-grained force field Martini (Souza and Marrink, 2020) and the program package Gromacs 2018.1 (Abraham et al., 2015). The protein was described using a Gõ-like model in combination with Martini (Poma et al., 2017; Souza et al., 2019). We used the tubbyCT crystal structure (Santagata et al., 2001) and modeled the missing loops with I-TASSER (Yang et al., 2015). To generate the CG model the missing loops were ligated to the crystal structure. This step was necessary to maintain the side chain orientations around the PI(4,5)P_2_ binding pocket of tubbyCT identified in the crystal structure. The potential depth of the Lennard-Jones potentials of the Gõ-like model was set to ε = 12 kJ/mol. More details about the protein model can be found in reference (Thallmair et al., 2020).

The simulation setup is schematically shown in Fig. 5I. The tubbyCT protein was placed in the water phase above a lipid bilayer consisting of POPC and a minor fraction of PI(4,5)P_2_. To study the competitive binding behavior of tubbyCT with increasing PI(4,5)P_2_ concentration, one leaflet contained 1 mol% of PI(4,5)P_2_ in all systems, while the other leaflet contained concentrations ranging from 1-5 mol% PI(4,5)P_2_. The system was neutralized, and solvated in a 0.15 M solution of NaCl. After the equilibration (500 ps), three replicas of each setup were simulated for 50 µs each. Thus, a total simulation time of 750 µs was acquired.

To analyze the binding behavior of tubbyCT, the distance between the PO_4_ plane of the bilayer and the PI(4,5)P_2_ binding pocket was analyzed. Integration of the population up to a PO_4_– pocket distance of 2.25 nm yielded the membrane-bound fraction of tubbyCT.

## ACKNOWLEDGMENTS

We like to thank Drs. Tobias Meyer, Lawrence Shapiro, Yasushi Okamura, Pietro de Camilli, Jen Liou, and Saikat Mukhopadhyay for providing expression plasmids and Drs. Christian Halaszovich and Thomas Berger for helpful discussions and critically reading the manuscript. S.T. thanks the Center for Information Technology of the University of Groningen for providing access to the Peregrine high performance computing cluster.

This work was supported by the DFG Research Training Group 2213 “Membrane Plasticity in Tissue Development and Remodeling.”

## AUTHOR CONTRIBUTIONS

Conceptualization, V.T., S.T., and D.O.; Investigation, V.T., L.S., S.E., S.T., C.G., and M.G.L.; Writing, D.O. and V.T.; Funding Acquisition, D.O.; Supervision, D.O..

## DECLARATION OF INTERESTS

The authors declare no competing interests.

---

**Supplemental Figure S1.**
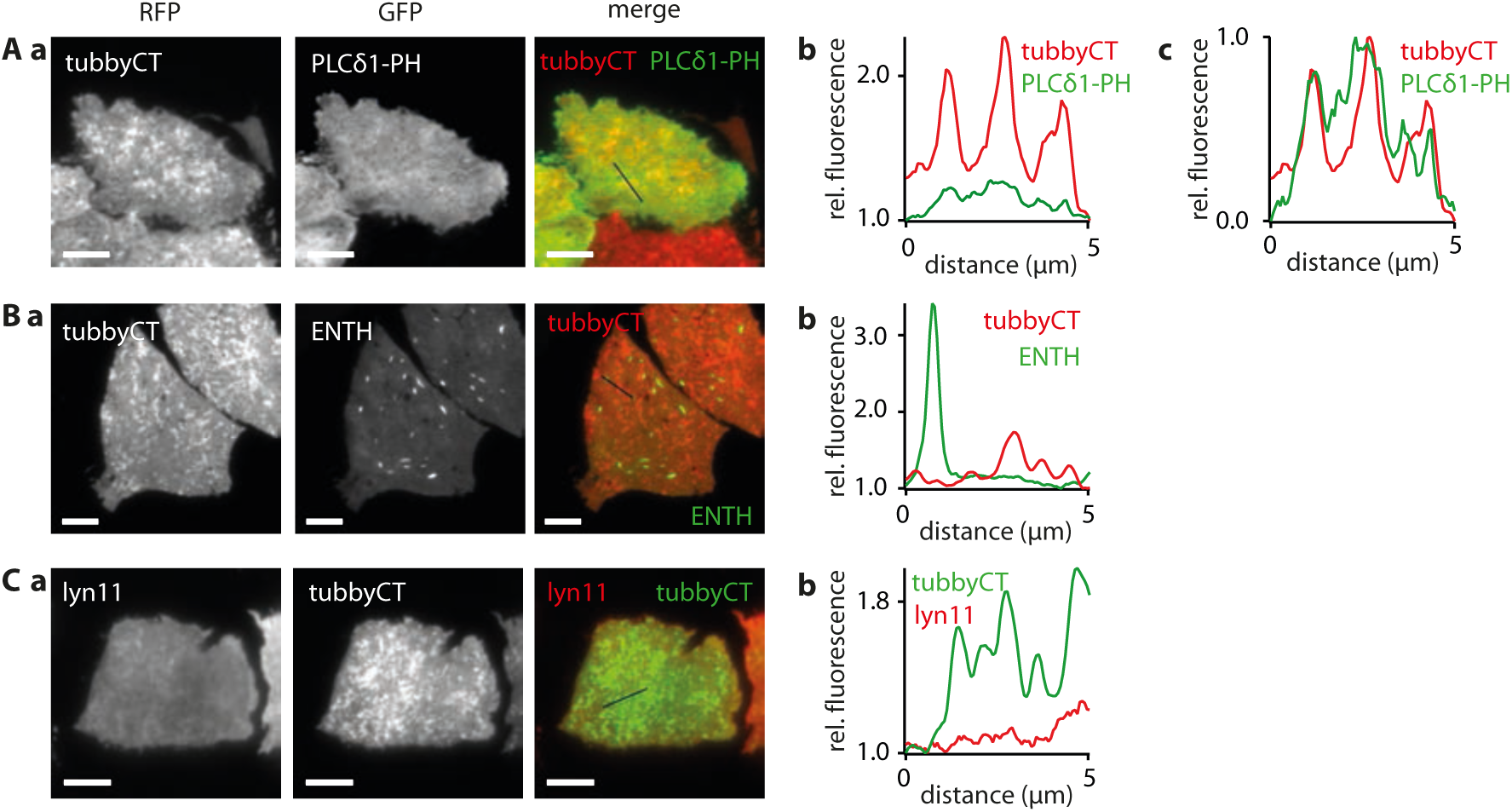
Lack of co-localization of tubbyCT clusters with other PI(4,5)P_2_ sensors and a non-specific membrane marker. **(A)a**, representative TIRF images of CHO cells co-transfected with RFP-tubbyCT and GFP-PLCδ1-PH indicates little enrichment of the PH domain in the clusters demarcated by tubbyCT. **b**, line profile plot as indicated in the merged fluorescence image. Fluorescence intensity normalized to minimum value along the profile. **c**, normalization of fluorescence signals to maximum intensity difference along the profile reveals a slightly higher abundance PLCδ1-PH in the tubbyCT cluster regions. **(B)** Co-localization analysis of tubbyCT co-expressed with PI(4,5)P_2_ sensor ENTH-GFP as in (A) shows lack of enrichment of the ENTH domain in tubbyCT clusters. **(C)** Lack of clustering of the general membrane marker lyn11-RFP and lack of co-localization with tubbyCT. TIRF imaging (a) and line profile analysis (b) as in (A,B). Scale bars, 5 µm.

**Supplemental Figure S2.**
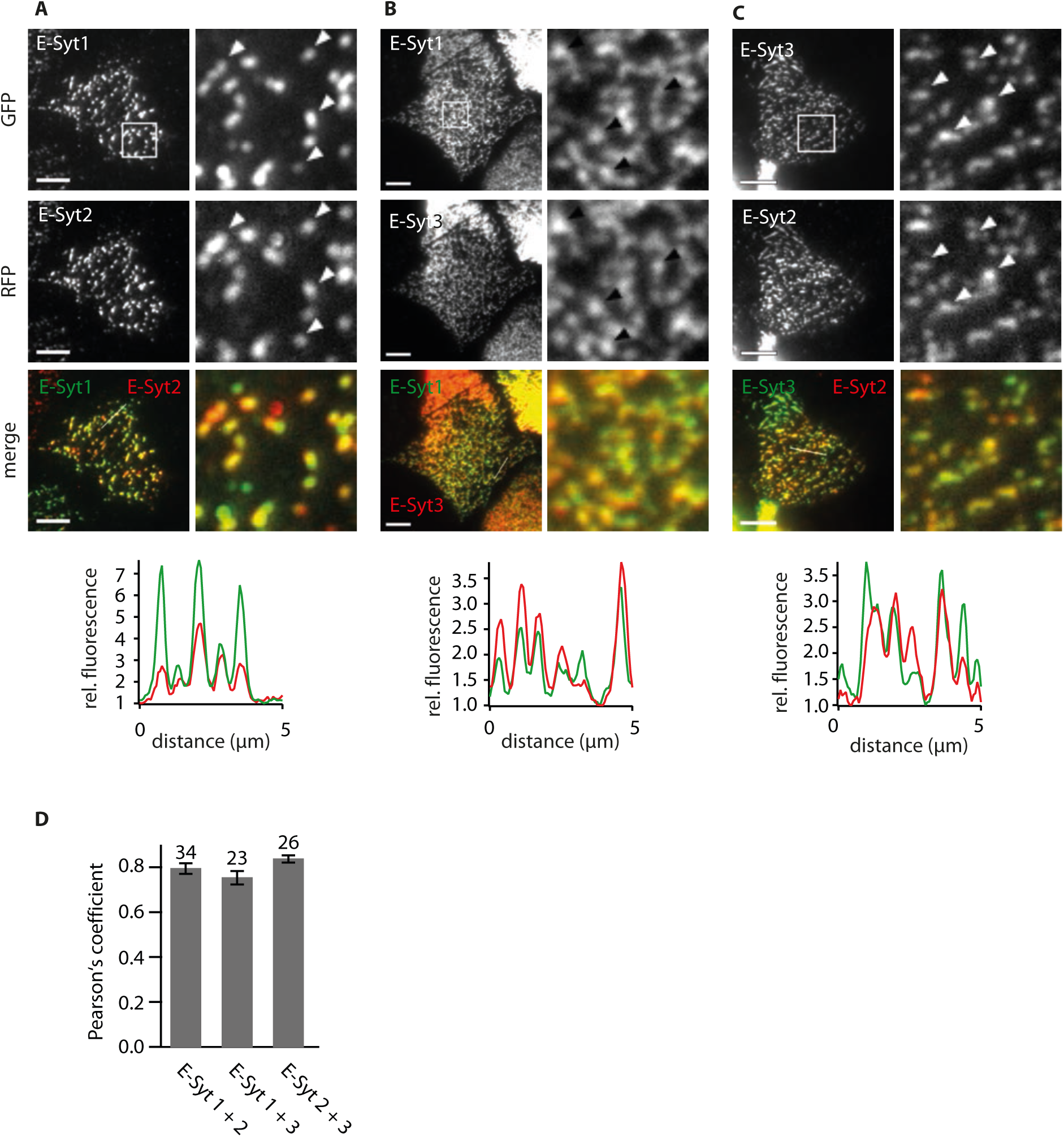
Co-localization of E-Syt isoforms. **(A-C)** Co-localization analysis of GFP-E-Syt1 co-expressed with RFP-E-Syt2 (A), GFP-E-Syt1 with RFP-E-Syt3 (B) and GFP-E-Syt3 with RFP-E-Syt2 (C). CHO cells were were transiently transfected with the respective plasmids and imaged by TIRF microscopy. Line profiles (length 5µm as indicated in merged images) are plotted below merged images. Fluorescence intensities are normalized to minimal values. In merged images and line profiles RFP-tagged proteins are displayed in red and GFP-tubbyCT in green. Scale bars, 5 µm. **(D)** Pearson’s coefficients (mean ± SEM) from cells as in (A-C); numbers of cells analyzed indicated.

**Supplemental Figure S3.**
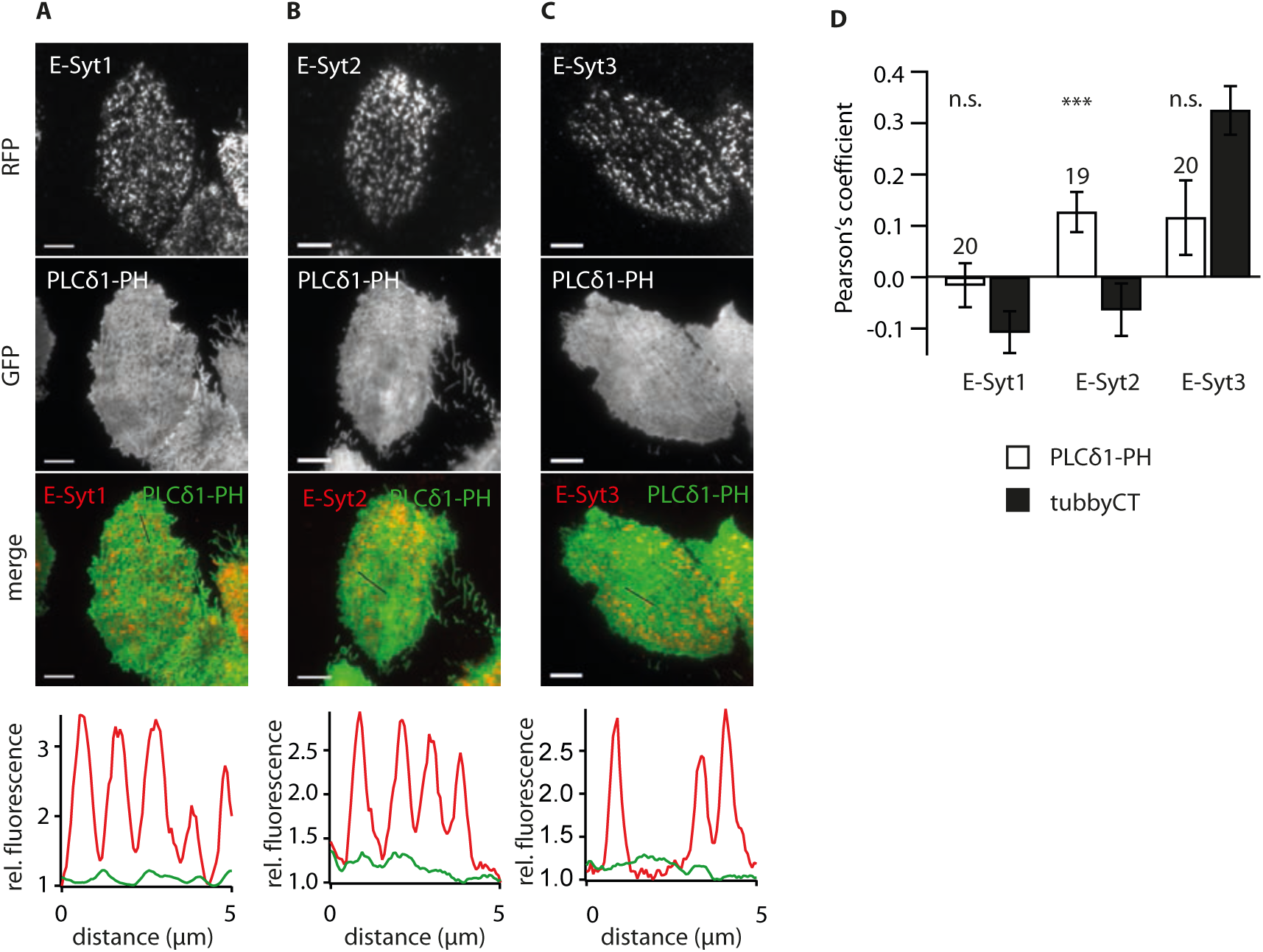
Analysis of co-localization of PLCδ1-PH-GFP with E-Syts. **(A-C)** Co-localization analysis of PLCδ1-PH-GFP with RFP-E-Syt1 (A), RFP-E-Syt2 (B) and RFP-E-Syt3 (C). CHO cells were transiently transfected with the respective plasmids and imaged by TIRF microscopy. Line profiles (5µm as indicated in merged images) are plotted in lower panels. Fluorescence intensities are normalized to minimal values. RFP fluorescence GFP fluorescence plotted in red and green, respectively. Scale bars, 5 µm. **(D)** Pearson coefficients r (mean ± SEM; number of cells indicated) were obtained from images as shown in (A-C). For comparison, data for GFP-tubbyCT colocalization are replotted from Figure 3f. (H0 = 0; n.s. = not significant; *: p < 0.05; ***: p < 0.01).

**Supplemental Figure S4:**
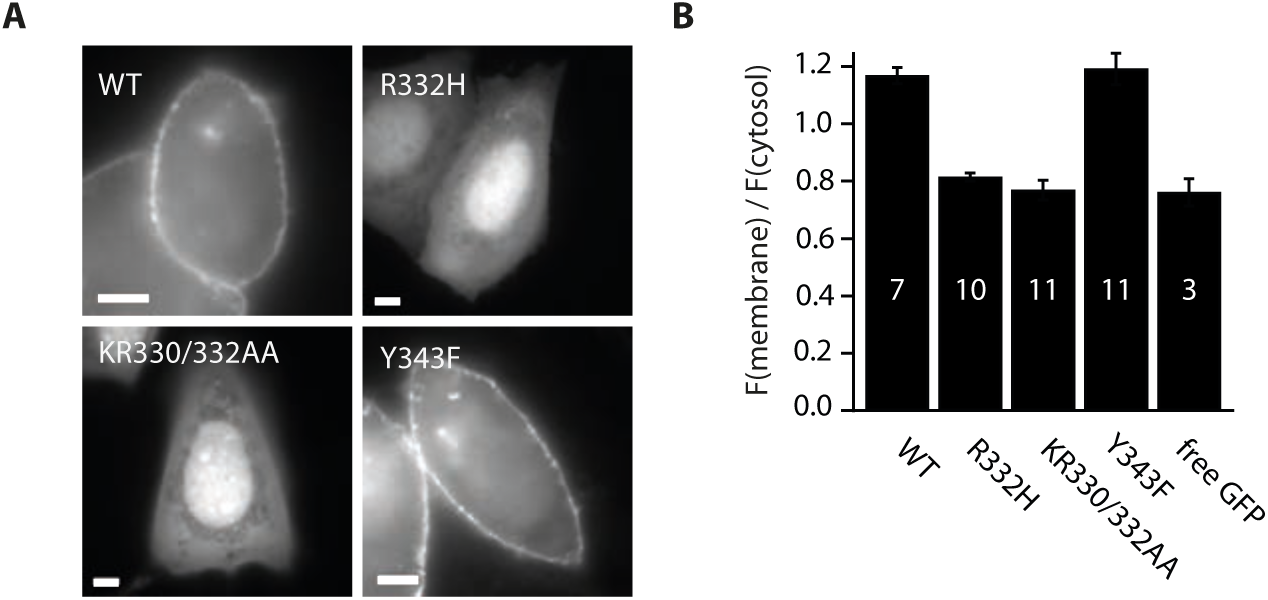
PI(4,5)P_2_ affinity of tubbyCT mutants estimated from membrane association. **(A)** Representative widefield fluorescence images of CHO cells expressing GFP-tubbyCT wild-type (WT) and mutants R332H, KR330/332AA, and Y343F. Note the different degrees of membrane association. Scale bars, 5 µm. **(B)** Ratio of membrane-to-cytosolic fluorescence of the tubbyCT mutants and free GFP for comparison (mean ± SEM; number of cells analysed as indicated). Ratios were determined from line profiles with localization of the membrane determined as the local fluorescence maximum of co-expressed membrane marker lyn11-RFP (not shown).

**Supplemental Figure S5.**
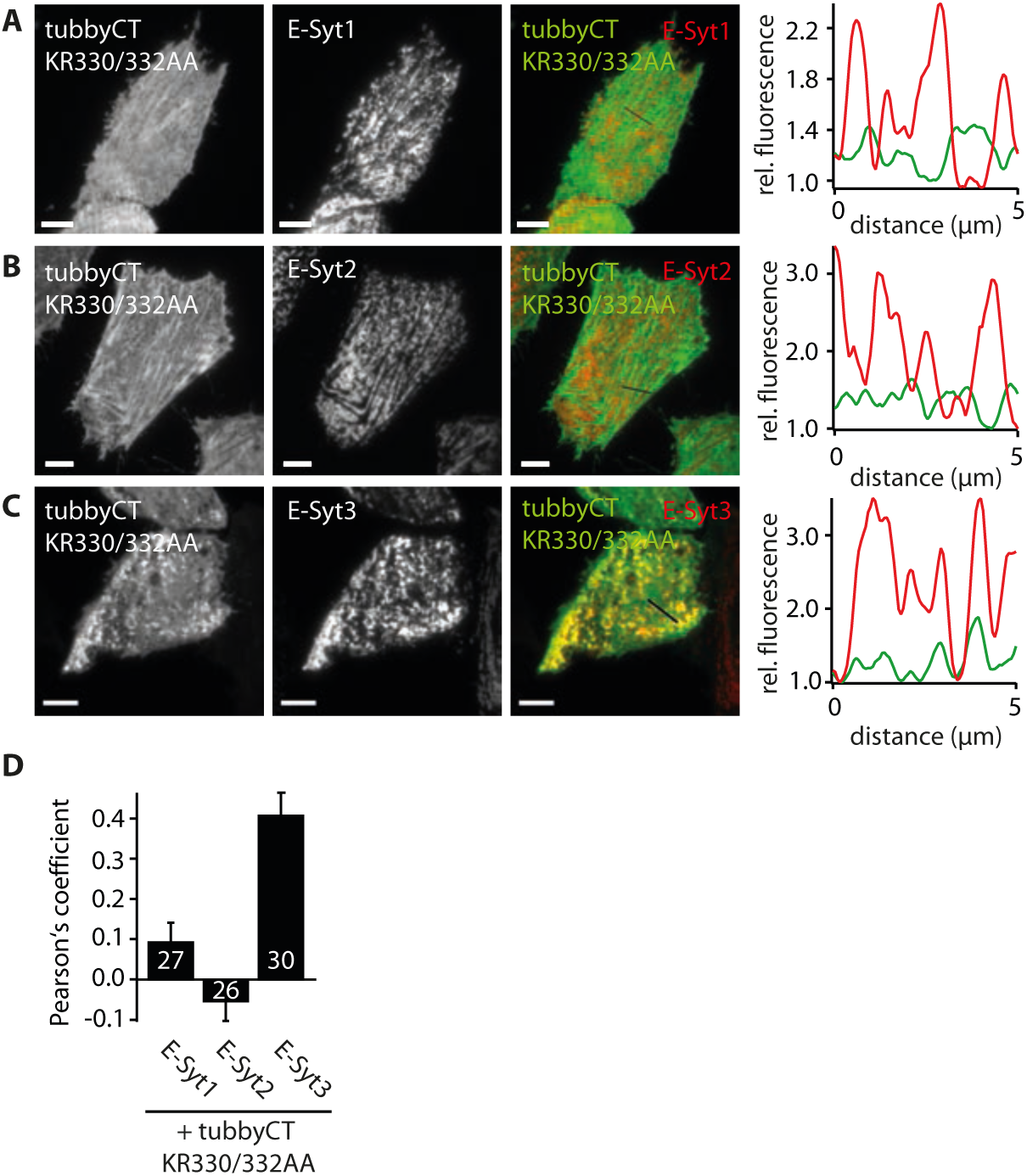
Over-expression of RFP-E-Syt3 but not RFP-E-Syt1/2 recruits GFP-tubbyCT KR330/332AA into ER-PM junctions. **(A-C)** Representative TIRF images of CHO cells co-expressing GFP-tubbyCT KR330/332AA and RFP-E-Syt1 (A), RFP-E-Syt2 (B) or RFP-E-Syt3 (C). Example cell in (C) is taken from Figure 6D. Scale bar = 5 µm. Line profiles (5µm; location highlighted in merged images) are plotted in the right panels qwith fluorescence intensities normalized to minimal values. In merged images and line profiles GFP-tubbyCT KR330/332AA is displayed in green and RFP-E-Syt in red. **(D)** Quantitative analysis of colocalization between GFP-tubbyCT KR330/332AA and the E-Syt isoforms indicated (mean ± SEM).

## Notes

### Competing Interest Statement

The authors have declared no competing interest.

